# A Unifying Mechanism for Synaptic Amyloid β Toxicity β Adrenergic Potentiation of the Ca^2+^ Channel CaV1.2 by Amyloid β

**DOI:** 10.64898/2026.04.25.720803

**Authors:** Peter Bartels, Sarah Rougé, James D. Scripter, Zhuoer Zeng, Zoila Maribel Estrada-Tobar, Jennifer Price, Ariel A. Jacobi, Ruben A. Berumen, Sheng-Yang Ho, Andranik Avedisyan, Yang K. Xiang, Chao-Yin Chen, Madeline Nieves-Cintron, Manuel F. Navedo, Mary C. Horne, Martin Hruska, Johannes W. Hell

**Affiliations:** Department of Pharmacology, University of California Davis, CA 95616-8636, USA; Department of Neuroscience, Rockefeller Neuroscience Institute, West Virginia University, Morgantown, WV 26506, USA; VA Northern California Health Care System, Mather, CA 95655, USA

**Author notes:** Corresponding authors: Peter Bartels, Mary Horne, Martin Hruska, Johannes W. Hell (Lead contact). These authors contributed equally to this work (Peter Bartels, Sarah Rougé).

## Abstract

Amyloid β peptides (Aβ) trigger Alzheimer’s disease (AD) but how has remained elusive. Aβ stimulates the β_2_ adrenergic receptor (β_2_AR), which forms a unique signaling complex with the L-type Ca^2+^ channel (LTCC) Ca_V_1.2. LTCCs have been implicated in the etiology of dementia and AD. We show that Aβ acutely potentiates Ca_V_1.2 via the β_2_AR, which triggers postsynaptic recruitment of Ca^2+^ permeable (CP) AMPARs in hippocampal cultures and impairs LTP in hippocampal slices within minutes. The long-term consequence is a loss of postsynaptic structure of glutamatergic synapses and neurotoxicity. Disrupting this signaling cascade with highly specific tools prevented all of these effects, unifying a number of currently divergent findings on Aβ synaptotoxicity including dysregulation of AMPARs and synaptic plasticity.

**TEASER:** Amyloid β peptide is the primary pathological agent in Alzheimer’s disease. It affects the nanoscale structure and function of glutamatergic synapses. The molecular mechanisms are largely unknown except for identification of several binding proteins including the β_2_ adrenergic receptor. We show that this binding potently (EC_50_<100 nM) augments Ca^2+^ influx through the L-type Ca channel Ca_V_1.2. This effect leads to improper recruitment of Ca^2+^-permeable glutamate receptors to postsynaptic sites (EC_50_<100 nM), synaptic dysfunction and ultimately neuronal death. This work identifies an essential mechanism in amyloid β neurotoxicity and explains many of the observed postsynaptic alterations.

**Highlights:** **Immediate effects of Aβ-induced stimulation of β_2_AR on Cav1.2:**

- Aβ induces phosphorylation of Cav1.2 on S1928 by PKA
- Aβ augments Cav1.2 activity via β_2_AR-induced S1928 phosphorylation within seconds

**Aβ-induced β_2_AR - Cav1.2 signaling has the following synaptotoxic effects.**

- Aβ induces postsynaptic accumulation of Ca-permeable AMPARs via β_2_AR - Cav1.2 signaling within 20 min
- Aβ impairs long-term potentiation (LTP) via β_2_AR - Cav1.2 signaling
- Aβ impairs postsynaptic structure and neuronal viability over 24 h
- Potency of Aβ in <u>all</u> the above effects is very high (100 nM Aβ is saturating!)
- <u>All</u> effects are prevented in S1928A KI mice and acute displaces β_2_AR from Cav1.2 with tat-Pep_1923_

## MAIN TEXT

Accumulation of Amyloid beta (Aβ) peptides is the defining hallmark of AD. Numerous reports show that Aβ exert deleterious effects on synaptic transmission and plasticity (*1–9*) as well as pericyte-mediated control of cerebral blood flow (*10, 11*). The precise molecular mechanisms underpinning the pathogenic effects of Aβ at postsynaptic sites are not well known. The long standing Ca^2+^ hypothesis posits that chronic elevation of Ca^2+^ influx via LTCCs causes neuronal dysfunction that underlies senile symptoms and AD. Although a number of studies support this hypothesis (*11–18*) it is unclear how Aβ potentiates LTCCs (but see (*18*)) and how this potentiation relates to synaptic dysfunction. In further support of a central role of Ca_V_1.2 dysfunction in brain pathologies are a large number of genome-wide association studies that identified Ca_V_1.2 gene variants as the overall most prominent risk factors for multiple brain disorders (*19–24*).

Aβ binds to the β_2_AR inducing cAMP production (*25*). Ca_V_1.2 forms a unique signaling complex with the β_2_AR, the trimeric Gs protein, adenylyl cyclase (AC), and the cAMP-dependent protein kinase PKA (*26, 27*). Association of these proteins with Ca_V_1.2 is required for potent upregulation of channel activity by β_2_AR signaling (*27–30*). The energetically costly formation of this complex suggests that Ca_V_1.2 is a major target for signaling by the β_2_AR. Collectively, these observations lead to the hypothesis that Aβ affects neuronal function at least in part through a β_2_AR - mediated increase in Ca_V_1.2 activity. Consistently, a putatively prominent role of the β_2_AR in AD is emerging (*31*). We now show that Aβ upregulates Ca_V_1.2 activity via channel-associated β_2_AR. This upregulation induces an increase in postsynaptic accumulation of AMPARs and especially of Ca^2+^-permeable (CP) AMPARs within minutes and impairs LTP. Ultimately, these effects result in a loss of synaptic nanostructure and neurotoxicity, which are prevented by inhibiting the β_2_AR and Ca_V_1.2.

## RESULTS

### Aβ augments Ca_V_1.2 activity via β_2_AR signaling

To test whether Aβ affects localized LTCC activity via β_2_AR signaling, cell-attached single channel recordings were performed on rat hippocampal cultures (HC) at 10-20 days in vitro (DIV) (Fig. 1A). This configuration provides a highly quantitative read-out for channel activity and affords the isolation of LTCC in the presence of blockers of non-LTCCs (see below) (*27–30*). As others (*32-^35^*),we used synthetic Aβ_1-42_ to prepare a mixture of Aβ monomers and oligomers. Inclusion of 10 μM total Aβ in the recording pipet augmented the channel availability, open probability (Po) of individual channels and peak current determined from the ensemble averages of all recordings (Fig. 1B,C). Analysis of the channel kinetics showed that the increase in Po by Aβ was due to a significant reduction in mean closed time (MCT) of individual channels, which globally led to an increase in mean total open time (TOT) through a higher number of unitary events per 100 sweeps (fig. S1A). However, no change was found in channel mean open time (MOT) of individual events (fig. S1A). Notably, the impact of Aβ on LTCC activity was prevented by the highly selective β_2_AR blocker ICI118551 but not the β_1_AR blocker CGP20712 (Fig. 1B, C, fig. S1A), implicating the β_2_AR in the increase in LTCC activity.

**Fig. 1.**
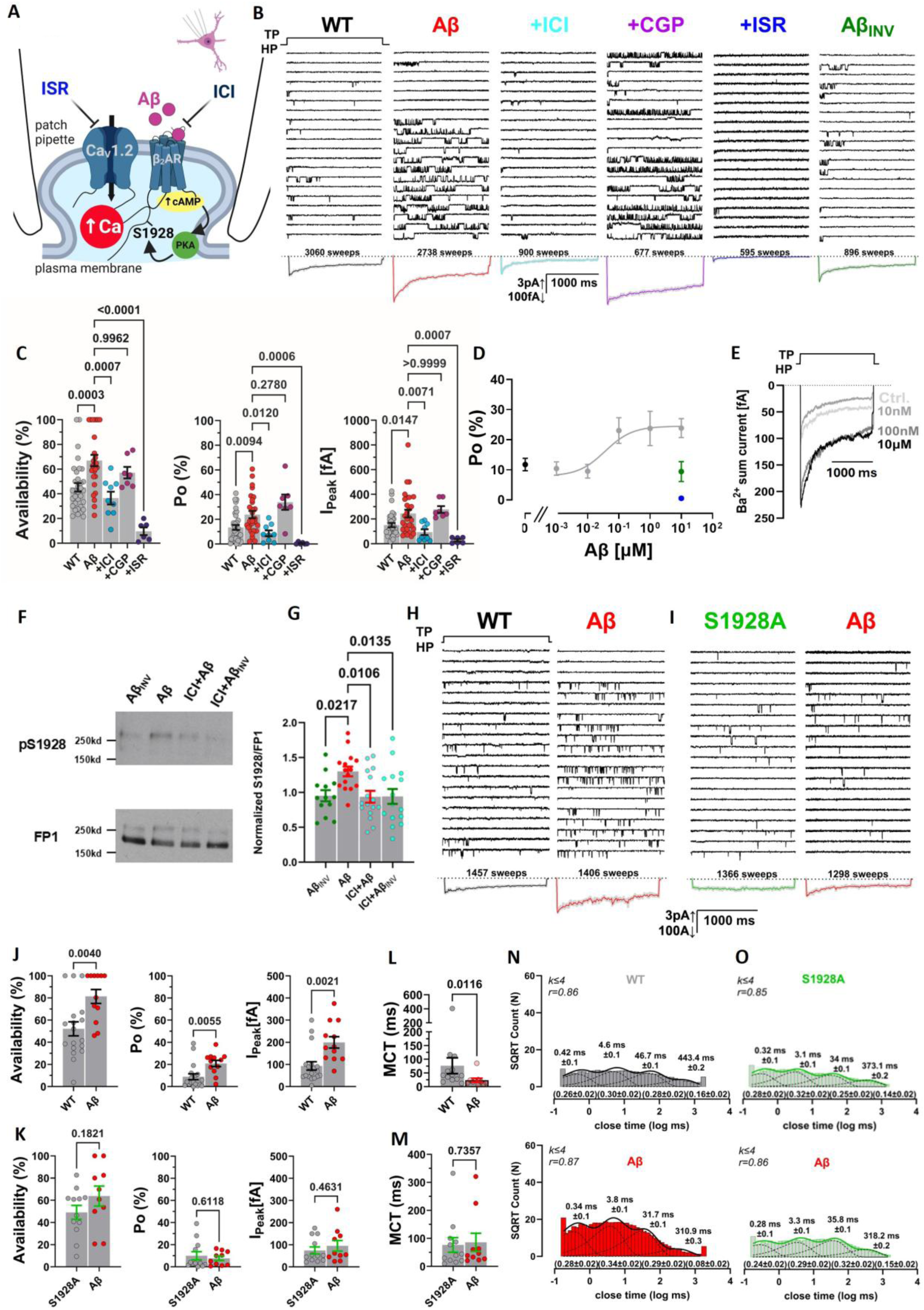
Aβ increased Ca_V_1.2 activity via β_2_AR-triggered S1928 phosphorylation. **A**: Configuration of cell-attached, single channel recordings of LTCC activity and conceptualization of Ca_V_1.2 upregulation by Aβ via the β_2_AR/Gs/AC/cAMP/PKA-S1928 pathway. All drugs including Aβ were present in the recording electrode only. **B**: Recordings of channel openings from rat primary HCs at 15-25 DIV upon depolarizations from a holding potential (HP) of -80 mV to a test potential (TP) of 0 mV for 2 s (depicted on top of traces for Control). Twenty consecutive exemplary, traces of LTCC activity for patches containing two channels (*k*=2). The pipette solution contained vehicle (WT) or Aβ either alone (Aβ, 10 µM) or in combination with the β_2_AR blocker ICI118,551 (ICI, 100 nM), the β_1_AR blocker CGP20712 (CGP, 100 nM), or the LTCC blocker Isradepine (ISR, 10 µM). Aβ_INV_ depicts inclusion of sequence-inverted inactive Aβ (10 µM). Bottom: mean ensemble average currents calculated from all recordings for each treatment. Scale bars indicate unitary current in pA (**↑**) and ensemble average current in fA (**↓**), respectively. **C**: Quantification of population data after correcting for the number of observed simultaneous openings (k≤4). The fraction of traces with LTCC activity over all traces (availability), the single channel open probability (Po), and the current maximum of the sum of each recorded data set (I_peak_) was increased by 10 µM Aβ, which was blocked by ICI but not CGP. Statistical significance was tested by a one-way ANOVA with Bonferroni post-hoc test, *p<0.05. **D**: Concentration dependence of the Aβ effect on Po. The calculated EC_50_ was 36.6 nM (95% CI: 6.4 nM - 363.2 nM). Green circle: Po recordings with 10 µM Aβ_INV_, which did not augment Po. Blue circle: Po recordings with 10 µM isradipine co-applied with 10 µM Aβ, which blocked virtually all currents. Data are represented as mean values ±SEM. A two-tailed unpaired T-test (titration) or a Welch-test for unequal variances were calculated for single comparison of each concentration or condition (Aβ_INV_ or ISR) against control data (**p<0.0042 and ****p<0.0001). **E**: Mean ensemble average data from all recordings in each group under control conditions (light grey) and Aβ at 10nM, 100nM and 10µM (darker grey to black) **F, G**: Forebrain slices were treated for 25 min with 1 µM Aβ or Aβ_INV_ either alone or with 100 nM ICI118551 before immunoprecipitation of Ca_V_1.2 and immunoblotting with anti-phospho S1928 and reprobing for total Ca_V_1.2 (**F**). Each dot in the bar diagram (**G**) represents the ratio of pS1928 to total Ca_V_1.2 signal for each semi-slice from 5 different mice. **H-O**: Recordings from mouse primary HCs at 15-20 DIV. The pipette solution contained vehicle (WT) or Aβ (10 µM). HP was -40 mV and TP 0 mV for 2 s. **H, I**: Twenty consecutive exemplary traces of LTCC activity from WT (left two columns) and S1928A KI neurons (right two columns). Comparative recordings exemplify one channel patches (*k*=1) with their mean ensemble average currents of all recordings from each group (below). **J, K**: Aβ augmented availability, Po, and I_PEAK_ in WT (**K**) but not S1928A KI HCs (**L**). Statistical significance was tested by an unpaired, two-tailed Student’s T-test, p<0.05%. **L, M**: Channel Mean closed time (MCT) under control condition and Aβ for WT and S1928A. **N, O**: Gamma distribution of closed state lifetimes for control and Aβ treatment for all recordings (pooled). Histograms were best described by the sum of four exponential close times (for details see Methods). **A**ll data are prepresented as mean ± SEM.

The nominal concentration of 10 μM Aβ is high and effects have been reported with as low as 100-200 nM Aβ (*32–35*). Therefore, we tested the effect of 1 nM to 1 μM Aβ on Po of LTCCs. There was no detectable effect by 1 and 10 nM Aβ but 100 nM Aβ showed a near maximal effect that was comparable to the increase in Po at 1 and 10 μM Aβ (Fig. 1D,E). The LTCC blocker isradipine nearly completely blocked all currents with 10 μM Aβ present (Fig. 1B, C), indicating that all currents including those due to the Aβ stimulation arose nearly exclusively from LTCCs. Control peptide with the inverse sequence of Aβ (Aβ_INV_) had no effect on Po even at 10 μM (green dot in Fig. 1D).

The β_2_AR upregulates Ca_V_1.2 activity via PKA-mediated phosphorylation of S1928 (*30*). Notably, treatment of forebrain slices with 1 μM Aβ for 20 min significantly augmented S1928 phosphorylation versus control treatment with 1 μM Aβ_INV_ (Fig. 1F, G). As observed for the Aβ-mediated increased in voltage-gated Ca^2+^ channel (VGCC) currents, this effect was completely blocked by 100 nM ICI118,551, which had no effect when co-applied with control Aβ_INV_. Thus, Aβ stimulates S1928 phosphorylation via β_2_AR signaling whose functional role was analyzed in S1928A KI mice. In WT mouse neurons 10 μM Aβ significantly increased the channel availability, Po, peak current of the ensemble averages, TOT, and frequency of unitary events of LTCC single channel currents and decreased MCT without affecting MOT (Fig. 1H,J,L; fig. S1B). Notably, none of these parameters were augmented in S1928A KI neurons (Fig. 1I,K,M; fig. S1C). These results unequivocally demonstrate that Aβ augments LTCCs activity by stimulating S1928 phosphorylation. They furthermore confirm that Ca_V_1.2 rather than Ca_V_1.3, which is the only other but much less prominent LTCC in the brain (*36, 37*), is the central effector for aberrant Ca^2+^ signaling induced by Aβ.

To compare the Aβ effects with those of classic βAR stimulation, we performed the same detailed analysis of gating kinetics for our earlier single channel recordings from HC (*30*). Consistent with signaling by Aβ via a βAR, the βAR agonist isoproterenol (ISO) increased channel availability, Po, and peak current (fig. S2A,B), as described earlier for the effects of ISO on Ca_V_1.2 in cardiomyocytes (*38*). These effects were completely abrogated by ICI (fig. S2C, D). The ISO-induced, ICI-sensitive increase in Po could be attributed to a reduction in MCT leading again to an overall higher TOT (fig. S2E, F).

To gain further mechanistic insight, we analyzed the channel state lifetimes by quantifying the distribution times of channel open and closed states. The underlying gamma distributions were overall best described with three open states and four closed states, using Maximum Likelihood (MLE) fitting. (Fig. 1N; fig. S3). Aβ showed minimal effects on the open states but significantly reduced the closed state duration, particularly affecting the slowest transition, leading to an overall higher opening frequency in WT neurons (Fig. 1N, fig. S3) but not S1928A KI neurons (Fig. 1O, fig. S4). This fine-grained analysis confirmed that Aβ significantly reduces Ca_V_1.2 closed time transitioning to subsequent channel openings resulting in a globally higher Po.

Direct binding of the β_2_AR to Ca_V_1.2 is required for upregulation channel activity by β_2_AR agonists (*29, 30*). We used the tat-tagged and therefore membrane-permeant synthetic peptide tat-Pep1923, which mimics the binding site of the β_2_AR within Ca_V_1.2 (residues 1923-1942 of the central, pore-forming α_1_1.2 subunit), to displace the β_2_AR from Ca_V_1.2 in HC and forebrain slices (*29, 30*). Acute application of tat-Pep1923, but not sequence-scrambled tat-Pep1923scr, completely reversed the Aβ-induced increase of LTCCs in channel availability, Po and peak current I_peak_ (Fig. 2A-D) and decrease in MCT (fig. S5A-D). Thus, the loss of Aβ effects on Ca_V_1.2 is not due to chronic changes in S1928A KI mice and requires the association of the β_2_AR with Ca_V_1.2 for the acute dysregulation of channel activity.

**Fig. 2.**
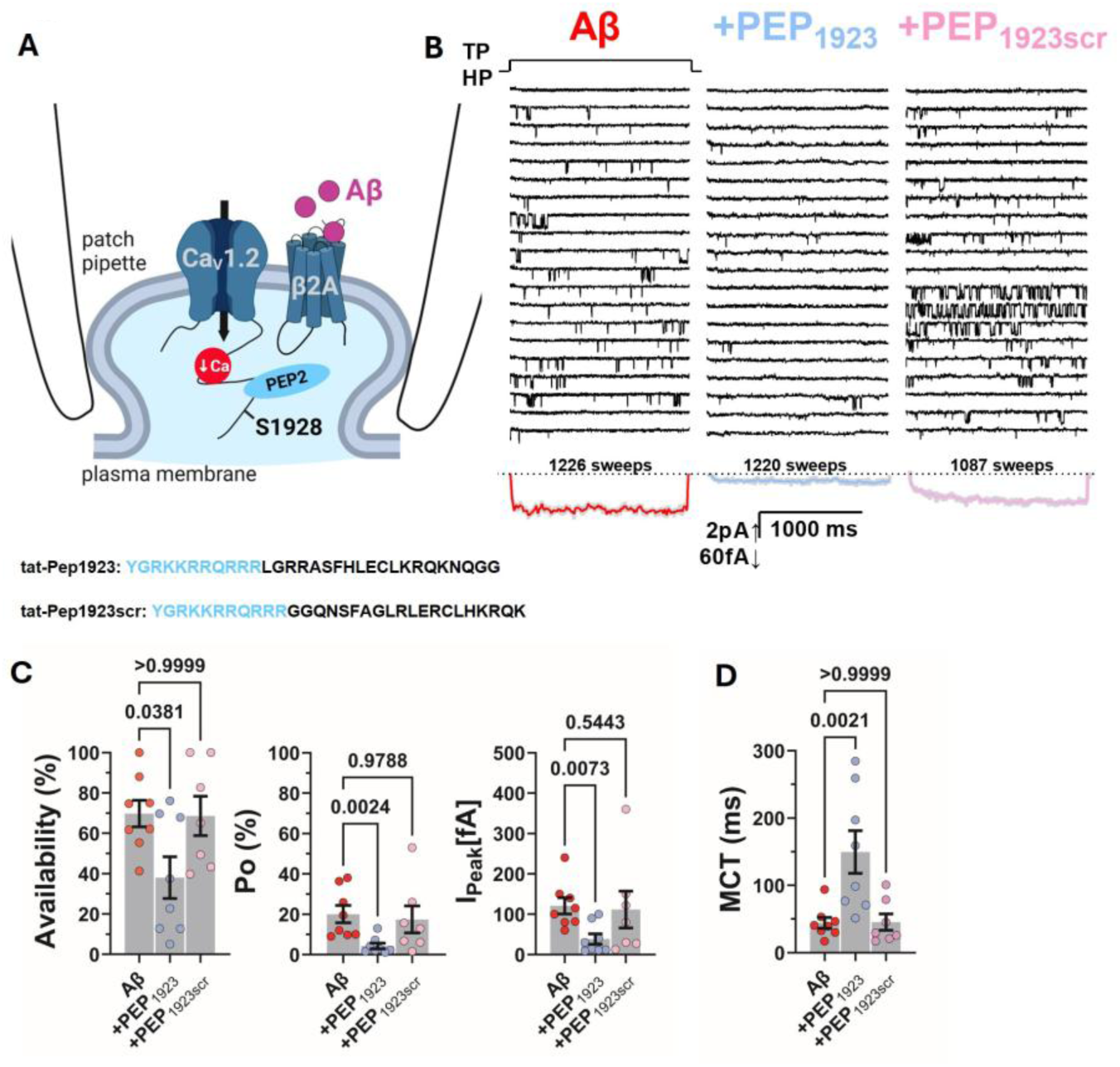
Acute displacement of the β_2_AR from Ca_V_1.2 prevented Aβ from upregulating channel activity. **A**: Conceptualization of displacement of the β_2_AR from its binding site on Ca_V_1.2 by Pep1923 (PEP2), which mimicks the binding site (*29*). Pep1923scr is a scrambled, inactive version of Pep1923. The tat sequence (cyan) makes the peptides membrane-permeant. **B-E**: Recordings from mouse primary HCs at 15-20 DIV of channel openings upop depolarizations from a holding potential (HP) of -40 mV to a test potential (TP) of 0 mV for 2 s (depicted on top of traces for Aβ alone). **B**: Twenty consecutive exemplary traces of patches containing two channels (*k*=2). The pipette solution contained 100 nM Aβ plus, if indicated, 10 µM tat-PEP1923 or tat-PEP1923scr vehicle. Bottom: mean ensemble average currents calculated from all recordings for each group. Scale bars indicate unitary current in pA (**↑**) and ensemble average current in fA (**↓**), respectively. **C**: Quantification of population data after correcting for the number of observed simultaneous openings (k≤4). The levels of availability, Po, and I_peak_ with Aβ alone were comparable to those in Fig. 1. tat-Pep1923 but not inactive tat-Pep1923scr dramatically reduced all three parameters to levels comparable to control conditions in Fig. 1. **D**: The low level of channel mean closed time (MCT) upon Aβ treatment was increased by tat-Pep1923 but not tat-Pep1923scr. Statistical significance was tested by a one-way ANOVA with Bonferroni post-hoc test, *p<0.05. All data: mean ± SEM.

To confirm the Aβ effects on Ca_V_1.2 activity in dendrites we performed Ca^2+^ imaging upon depolarization of HC with 90 mM K^+^. Application of 100 nM Aβ significantly augmented Ca^2+^ transients in WT but not S1928A KI HCs (fig. S6). This finding indicates that the β_2_AR – S1928 axis is prominent in dendrites and again identifies Ca_V_1.2 versus Ca_V_1.3 as the relevant Aβ target downstream of the β_2_AR.

### Aβ affects post-synaptic AMPAR composition via β_2_AR – Ca_V_1.2 signaling

AMPARs at cortical synapses are mostly GluA2 subunit-containing and hence mostly Ca^2+^ impermeable AMPARs (CI-AMPARs) (*39, 40*) (but see (*41, 42*)). However, pathological conditions can cause long-lasting postsynaptic insertion of Ca^2+^ permeable AMPAR (CP-AMPARs), which is thought to be detrimental to neuronal functions due to the chronically increased Ca^2+^ influx (*43–47*). As Aβ induces postsynaptic recruitment of CP-AMPARs (*34*) and LTCC activity stimulates the postsynaptic recruitment of GluA1 homomers (*48*), the prevalent CP-AMPARs in cortex (*49*), we hypothesized that Aβ induces postsynaptic CP-AMPAR accumulation by stimulating β_2_AR - Ca_V_1.2 activity.

To directly observe the impact of Aβ on the localization of CP-AMPARs, we undertook Stimulated Emission Depletion (STED) microscopy. GFP-transfected cortical neurons at 18-21 DIV were surface stained with fluorescently-labeled nanobodies against GluA1 and GluA2 (*50–52*), followed by fixation, permeabilization, and labeling of PSD-95 (Fig. 3A,B) (*53*). In vehicle-treated neurons most postsynaptic AMPAR nanoclusters as defined by colocalization with PSD-95 contained both, GluA1 and GluA2 (Fig. 3C), likely reflective of GluA1/GluA2 diheterotetramers (*40, 50*). PSD-95 nanodomains that contained only GluA2 and lacked GluA1 (“GluA2 only”) were also prominent (Fig. 3D), likely reflective of GluA2/GluA3 diheterotetramers (*40, 50, 54*). PSD-95 nanodomains containing GluA1 only, likely GluA1 homomers (*40, 49*), were rare (Fig. 3E). Treatment with 100 nM Aβ for 20 min increased GluA1 only PSD-95 nanoclusters (Fig. 3D, E). It decreased GluA2 only PSD-95 nanoclusters, which can be explained by the reduction in synaptic GluA2/GluA3 heteromers by Aβ (*55*). In contrast, no change was observed for colocalized GluA1/GluA2 nanoclusters. Importantly, blocking either β_2_AR or LTCC by pre-treating neurons with 400 nM ICI118551 or 10 µM isradipine, respectively, blocked the effects of 20 min exposure to 100 nM Aβ on GluA1 and GluA2 colocalization with PSD-95 (Fig. 3 D,E). These data indicate that 100 nM Aβ leads to changes in AMPARs composition at synapses within minutes and are consistent with the accumulation of GluA2-lacking CP-AMPARs to synaptic sites in a manner that requires β_2_AR and LTCC activity.

**Fig. 3:**
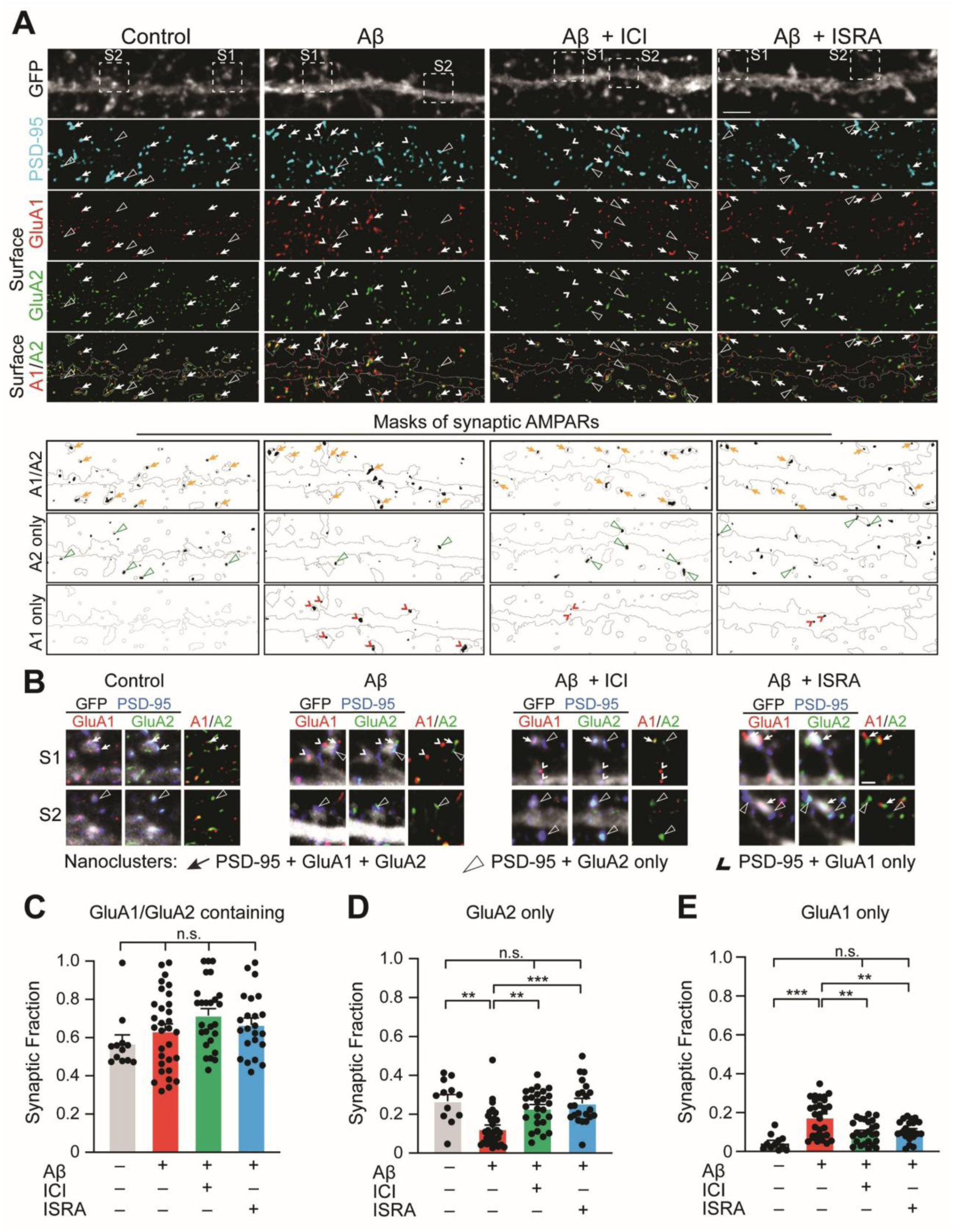
Aβ led to the accumulation of GluA2-lacking AMPARs at synapses via activation of β_2_ARs and LTCCs. **A**: Representative tau-STED images of surface-labeled GluA1 (red) and GluA2 (green) AMPAR subunits at synaptic sites identified by the staining for PSD-95 (blue) in DIV18-21 cortical neurons. Neurons were treated with vehicle or 100 nM Aβ for 20 minutes in the presence or absence of ICI118551 (ICI, 400 nM) or Isradipine (ISRA, 10 µM) before surface labeling. Imigang was confocal for GFP (gray) and STED for GluA1, GluA2 and PSD-95. Masks below indicate colocalized GluA1 and GluA2 (A1/A2, arrows), GluA1-lacking (A2 only, open arrowheads) and GluA2-lacking (A1 only, open arrowheads) nanoclusters at synapses based on colocalization with PSD-95. Scale bar: 2µm. **B**: High resolution of PSD-95 (blue), GluA1 (red) and GluA2 (green) in individual spines (GFP, gray; from A, white squares). Scale bar: 500 nm. **C-E**: Fraction of synaptic PSD-95 nanoclusters containing GluA1 and GluA2, GluA2 only, or GluA1 only (mean ± SEM; n = 12 neurons for control, 31 for Aβ, 26 for Aβ+ICI, 23 for Aβ+ISRA; ***p<0.0001, **p < 0.002, one way ANOVA with Tukey’s post-hoc). All data: mean ± SEM from three independent experiments.

In parallel, we monitored surface localization of GluA1 in forebrain slices from WT and S1928A KI mice following a 20 min treatment with 1 μM of either Aβ or Aβ_INV_ as control by surface biotinylation. Aβ significantly increased surface GluA1 content in WT but not S1928A KI mice versus Aβ_INV_ (fig. S7).

To examine functional consequences of this re-distribution of AMPAR subtypes, HC were treated at 14-20 DIV with 1 μM Aβ for 20 min prior to recording of miniature excitatory postsynaptic currents (mEPSCs; Fig. 4A-C). Aβ increased mEPSC amplitude by about 60% versus control treatment from about 25 pA to 40 pA (Fig. 4D). Subsequent application of IEM1460 after the initial 5 min recording of mEPSCs completely reverted the increase in mEPSC amplitude over the following 15 min (Fig. 4D). This time course is consistent with the use dependent effect of this blocker of CP-AMPARs as a channel must open for IEM1460 to block it during subsequent events. Aβ also decreased the time constant tau for the decay of the mEPSCs (Fig. 4C (middle), E). This finding is also consistent with recruitment of CP-AMPARs because their decay tau is thought to be in general faster than CI-AMPARs (*56–58*).

**Fig. 4.**
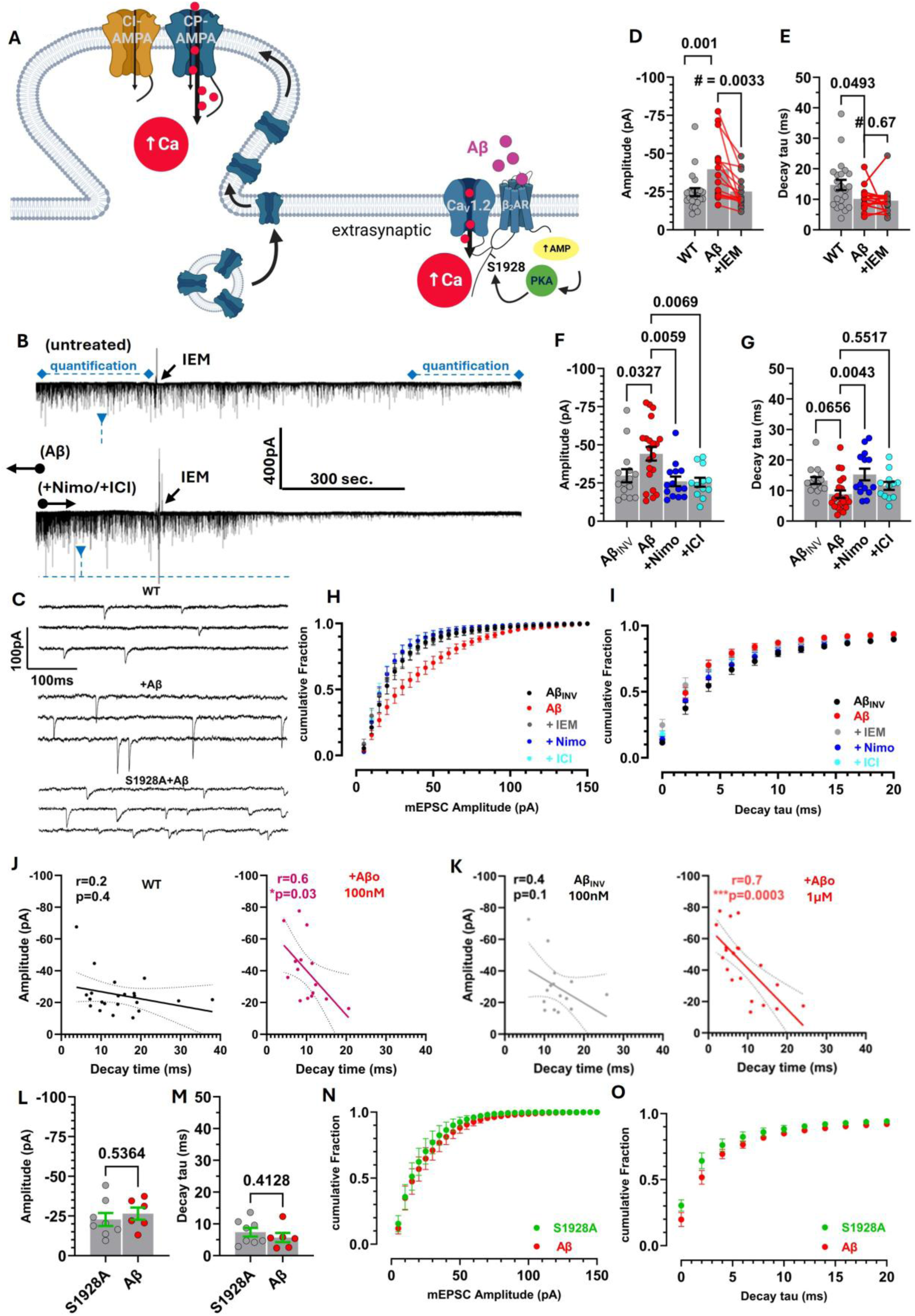
Aβ triggered postsynaptic recruitment of CP-AMPAR by stimulating β_2_AR and Ca_V_1.2 activity. **A:** Whole-cell patch recording of mEPSCs and conceptualization of postsynaptic recruitment of CP-AMPARs by increased Ca^2+^ influx via Aβ/β_2_AR/Gs/AC/cAMP/PKA/Ca_V_1.2 signaling. Ca^2+^ influx drives surface expression and postsynaptic accumulation of GluA1 homomers via unknown mechanisms (*48*). Mouse HCs were preincubated at 15-20 DIV with indicated compounds for 20-40 min before application of 1 µM TTX plus 50 µM picrotoxin, seal formation and recording of spontaneous mEPSCs at -60 mV. **B, C:** Representative traces of mEPSCs without and with preincubation of 1 µM (**B**) Aβ. Five min. after recording started, 50 µM IEM1460 was added and recording continued for further 15 min to allow the use-dependent block of CP-AMPARs by IEM1460 to fully develop. **D, E:** 1 µM Aβ significantly increased mEPSC amplitude and accelerated decay tau. IEM1460 reversed the increase in amplitude as measured 15-20 min after its addition. **F-I:** Population data (**F, G**) and cumulative fraction plots (**H, I**) of mEPSC amplitude and decay tau upon treatment with 100 nM inactive Aβ_INV_ as control or 100 nM Aβ, which increased mEPSC amplitude and accelerated decay tau to the same degree as 1 µM Aβ. Both effects are prevented by 10 µM nimodipine (+Nimo) and 400 nM ICI118551 (+ICI) when co-applied with Aβ. **J, K:** Correlation analysis of mEPSC amplitude versus decay tau. Treatment with 1 µM as well as 100 nM Aβ resulted in a steep relationship between amplitude and tau (r > 0.5), which was absent in control treatments with vehicle and Aβ_INV_, respectively. Pearson correlation was used to depict the relationship between amplitude and decay time, *p<0.05. **L-O:** Population data (**L, M**) and cumulative fraction plots (**N, O**) of mEPSC amplitude and decay tau upon treatment of S1928A KI HC with vehicle as control or 100 nM Aβ, which did not affect either parameter in these HCs. All data: mean ± SEM (p values: unpaired T-test or for > 2 groups one-way ANOVA with Bonferroni correction; #<0.05, paired t-test).

After establishing a robust effect of 1 μM Aβ on mEPSCs and its reversal by IEM1460, we tested 100 nM Aβ, which gave a near maximal effect on LTCC Po (Fig. 1D). Consistently, the increase in mEPSC amplitudes (∼ 40 pA) and decrease in decay taus (∼8 ms; Fig. 4F,G) seen with 100 nM Aβ were comparable to those for 1 μM Aβ (Fig. 4D,E). Accordingly, 100 nM Aβ was close to if not at saturation with respect to its effects on both, LTCCs and mEPSC. As negative control we used Aβ_INV_, which mirrored the above control amplitudes (∼ 25 pA) and decay taus (∼14 ms; Fig. 4F,G). Binned, cumulative fraction plots indicate that Aβ effects are fairly evenly distributed across the whole spectrum of mEPSC amplitudes and decay taus (Fig. 4H,I). Of note, the increase in amplitude and decrease in decay tau were completely prevented by the LTCC blocker nimodipine and the β_2_ AR blocker ICI118551 (Fig. 4F-I).

The decrease in the decay tau averages was strongly correlated with the increase in amplitude averages only after either 100 nM or 1 μM Aβ treatments but not under control conditions with vehicle or Aβ_INV_ (Fig. 4J, K). This effect on this correlation was analogous to that seen upon chronic neuronal inactivity, which also leads to postsynaptic recruitment of pharmacologically identified CP-AMPARs and the appearance of such a correlation between decay tau and amplitude of the mEPSCs (*57, 58*). Accordingly, neurons that showed minimal if any response to Aβ with respect to mEPSC amplitude averages also retained slow decay tau averages, whereas neurons that exhibited a strong increase in mEPSC amplitude acquired much faster decay tau averages. This correlation further supports the notion that the overall increase in mEPSC amplitude by Aβ was mediated by recruitment of CP-AMPARs with fast decay (*56–58*). However, AMPAR auxiliary subunits also affect decay tau (*47, 59*) including TARPs (*60*), CNIHs (*61, 62*), and SynDIG4 (*63*) and a change in decay tau could also reflect a change in auxiliary subunits. Importantly, Aβ did not at all affect mEPSCs amplitude or decay tau in S1928A KI mice (Fig 4L-O). Given that the Aβ effects were blunted by ICI118551, nimodipine, and in S1928A KI neurons, we conclude that acute Aβ treatment triggered recruitment of CP-AMPARs to postsynaptic sites by stimulating the β_2_ AR and thereby Cav1.2 activity.

### Aβ affect network functionality via β_2_AR - Ca_V_1.2 signaling

We used acute hippocampal slices to test the role of β_2_AR - Ca_V_1.2 signaling in Aβ toxicity in an intact neuronal network. Slice treatment with Aβ for 30-60 min impairs various forms of LTP and especially LTP induced by two 100 Hz / 1 s tetani that are 10-20 s apart (2x100 Hz LTP) (*5, 6, 8, 35, 64, 65*). Field excitatory postsynaptic potentials (fEPSPs) were recorded from CA1. Pre-treatment of the slices for 20 min with 1 μM Aβ nearly completely abrogated 2x100 Hz LTP (Fig. 5A, B). This effect was completely blocked by 10 μM nimodipine, implicating an important role of LTCCs in this effect. LTP was not affected by pre-treatment with 1 μM Aβ_INV_. We reduced the Aβ concentration to 200 nM to better match our conditions in HCs where 100 nM had a near maximal effect. Again, LTP was largely abrogated (Fig. 5C, D). Strikingly, Aβ did not at all impair LTP in S1928A KI mice, specifically implicating this phosphorylation site and thereby specifically Ca_V_1.2 (Fig. 5E, F).

**Fig 5.**
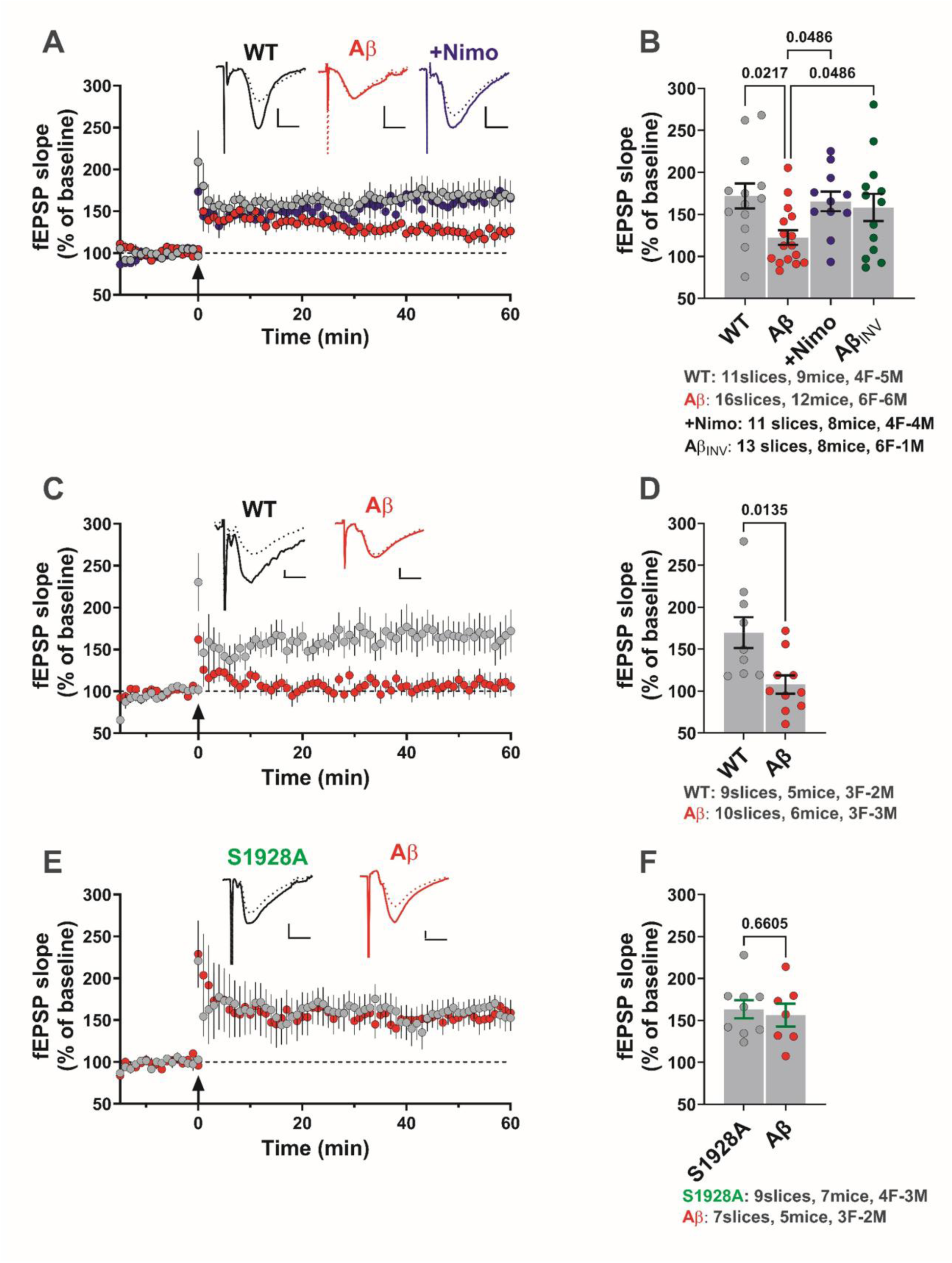
Aβ impaired LTP by engaging phosphorylation of Ca_V_1.2 on S1928. Field EPSPs were recorded from WT and S1928A KI hippocampi. LTP was induced by two 100Hz / 1 s tetani, separated by 10 s. Shown are time courses of initial fEPSP slopes (**A, C, E**) and population data of changes in synaptic transmission determined as the averages of fEPSP initial slopes 50-60 min after versus 0-10 min before tetani (**B, D, F**). Data are given as means ± SEM. *Inserts in A, C, E:* Sample traces of fEPSP recordings 10 min prior to tetanus (dotted lines) and 60 min post tetanus (solid lines). **A, B:** Wild-type slices were pre-treated for 20 min with vehicle (0.0001% DMSO; WT), 1 µM Aβ alone, 1 µM Aβ_INV_, or 1 µM Aβ plus 10 µM nimodipine (+Nimo). Aβ, but not Aβ_INV_ significantly impaired LTP, which was prevented by co-application of nimodipine. **C, D:** Wild-type slices were pre-treated for 1 h with vehicle (0.0001% DMSO) or 200 nM Aβ, which nearly completely abrogated LTP. **E, F:** S1928A slices were pre-treated for 1 h with vehicle (0.0001% DMSO) or 200 nM Aβ, which did not affect LTP in S1928A KI slices. All data: mean ± SEM (p values: B: one-way ANOVA with Holm-Šίdak correction; D, E: unpaired t-test).

### Effects of chronic Aβ treatment on postsynaptic nano-architecture and its cytotoxicity requires β_2_AR - Ca_V_1.2 signaling

To determine the role of β_2_AR - Ca_V_1.2 signaling in long-term effects of Aβ on synaptic integrity, we examined changes in PSD-95, the key organizer of excitatory postsynaptic sites (*66–69*), within dendritic spines using STED microscopy (fig. S8A)(*70–72*). Treatment of cortical cultures at 18-24 DIV with 500 nM Aβ for 24 h led to a reduction in density of dendritic spines, the postsynaptic sites of glutamatergic synapses in cortex and hippocampus (fig. S8B), consistent with previous results (*35, 73–75*). In the remaining spines (>60% of control) the fraction of those lacking a PSD-95 nanocluster increased and of those containing only one PSD-95 nanocluster decreased (fig. S8C,D). At the same time, there was an overall increase in the average number of PSD-95 nanoclusters per spine (fig. S8E). This effect reflects earlier findings that a high level of PSD-95 as present in spines with multiple nanodomains protects spines against damage by Aβ (*75*). All of the Aβ effects on postsynaptic nano-architecture were, once more, prevented by pretreating neurons with the β_2_AR antagonist ICI118551 or the LTCC blocker isradipine.

The number of PSD-95 nanoclusters correlates with spine size and thereby synaptic strength (*70, 71, 76, 77*). We investigated the effects of 24-hr Aβ treatment on spine morphology. While we did not find any differences in sizes of spines lacking PSD-95 nanodomains that were present after Aβ treatment, remaining spines with a single PSD-95 nanodomain were significantly larger (fig. S8F, G). Surviving single nanodomain spines may have started out as stronger synapses than those that lost PSD-95 upon Aβ application; perhaps PSD-95 content was higher in the former than latter spines, consistent with the protective effects of PSD-95 in Aβ synaptotoxicity (*75*). ICI118551 and isradipine also rescued the effects of Aβ on spine morphology, once more implicating β_2_AR and Cav1.2.

To specifically block β_2_AR-induced S1928 phosphorylation we utilized tat-Pep1923. Tat-Pep1923, but not inactive tat-Pep1923scr, prevented the Aβ-mediated decrease in overall spine density and number of spines containing a single PSD-95 nanodomain and the increase in spines lacking PSD-95 (fig. S9A-E). As before, Aβ did not reduce the proportion of spines with multiple PSD-95 nanodomains. These data support the idea that the effects of 100 nM Aβ are primarily limited to spines with initially low nanoscale complexity. Furthermore, spines with a single PSD-95 nanodomain that retained PSD-95 at the end of 24 hour period were larger with control tat-Pep1923scr when spines without a PSD-95 nanodomain were not (fig. S9F,G). Again, surviving spines might have started out with a higher PSD-95 content in their single nanodomains than spines that got eliminated. Overall, these findings suggest that β_2_AR-Ca_V_1.2 signaling mediates the chronic effects of Aβ, disrupting spine integrity and inducing changes in sub-structure.

Finally, we tested whether Aβ-induced cytotoxicity could be alleviated by blocking β_2_AR-Ca_V_1.2 signaling. Treatment of 11-13 DIV HCs with 1 µM Aβ for 48 h increased the portion of neurons with severe plasma membrane damage from 10 to 20% (fig. S10). This increase was completely prevented by isradipine, ICI118551, and tat-Pep1923.

## DISCUSSION

We identify upregulation of Ca_V_1.2 activity as a very early effect of Aβ on neuronal functions that occurred within seconds of Aβ exposure and triggered within minutes, an increase in the activity of postsynaptic CP-AMPAR in cultured neurons and a decrease in LTP at the network level that relied on phosphorylation of Ca_V_1.2 at S1928. These results build on our earlier work showing that aged rats have a significant increase in PKA-mediated phosphorylation of Ca_V_1.2 on S1928 (*18*). S1928 phosphorylation augments Ca_V_1.2 channel activity (*30*) and explains the increase in LTCC activity and related senile impairments in aging rodents shown by others (*12, 17, 78*). Notably, here we found that Aβ-mediated upregulation of Ca_V_1.2 activity was also responsible for changes in synaptic nanoarchitecture and neuronal viability that occurred over a 24-48 h period. We conclude that Aβ exerts its effects on synaptic functions and neuronal integrity to a large degree via upregulation of Ca_V_1.2 activity by β_2_ AR signaling and the ensuing phosphorylation of Ca_V_1.2 on S1928.

Although the Aβ - induced effects on Ca_V_1.2 activity and synaptic architecture and functions were completely reversed in S1928A KI mice or tat-Pep1923, it is well established that Aβ acts on other targets including mGluR1 (*35*) and mGluR5 (*2–4*), which can augment LTD (*79, 80*); PirB/LilrB2 (*32, 81*), thereby promoting synapse shrinkage through calcineurin-mediated cofilin activation; complement C1q (*33*), which fosters synapse elimination; and EphB2 (*82*) whose Aβ-induced impairment can trigger NMDAR removal. The complete reversal of all Aβ - induced effects in S1928A KI mice strongly implicates Ca_V_1.2 in Aβ – induced effects. The most parsimonious explanation is that dysregulation of Ca_V_1.2 is an essential component of the synaptic Aβ effects that has to occur in parallel or in series with the Aβ effects on mGluR1/5, PirB/LilrB2, C1q, EphB2 or other Aβ targets.

As an early event, S1928 phosphorylation induced by Aβ - β_2_ AR signaling constitutes a potential target for early pharmacological intervention in AD with some evidence implicating beta blockers in reducing risk for AD (*31*). At the same time, Ca_V_1.2 has numerous physiological functions in the brain, the cardiovascular system and beyond, increasing the likelihood of side effects of general LTCC blockers. Thus, we need to find alternative treatments that can specifically control pathological upregulation of Ca_V_1.2 in the brain. Our findings identify Aβ-β_2_AR-Ca_V_1.2 signaling and in particular the β_2_AR-Ca_V_1.2 interaction as a potential target for early pharmacological intervention in AD (*83, 84*). Given the increasingly sophisticated detection of early markers for AD (*85–90*), intervention such as disrupting Aβ-β_2_AR-Ca_V_1.2 signaling during early stages of AD is becoming increasingly conceivable. Importantly, despite the critical role of S1928 in upregulation of Ca_V_1.2 activity in neurons (Fig. 1) (*30, 84*), S1928 plays only a minimal role in regulation of Ca_V_1.2 in the heart where phosphorylation of the small G protein rad, which is not detectable in neurons, mediate upregulation of Ca_V_1.2 (*84, 91*). Thus, specifically inhibiting S1928 phosphorylation will most likely not affect cardiac function.

## MATERIALS AND METHODS

### Aβ and other Reagents

Aβ_1-42_ (Aβ; DAEFRHDSGYEVHHQKLVFFAEDVGSNKGAIIGLMVGGVVIA) and inverted Aβ_42-1_ (Aβ_INV_; AIVVGGVMLGIIAGKNSGVDEAFFVLKQHHVEYGSDHRFEAD) were from Royobiotech, Shanghai, China. Aβ and Aβ_INV_ were solubilized at a concentration of 1 mM in 0.1% DMSO. The solutions were vortexed for 15 seconds at 15-second intervals over a total period of 30 minutes. Aliquots were prepared on ice and stored at -80°C for no longer than one month. Each batch of reconstituted Aβ was validated for the presence of monomers, dimers, and trimers by running samples on Tricine SDS-PAGE gels, transferring them to a 0.2 µm PVDF membrane, and performing immunoblotting using anti-mouse Aβ antibody (1µg/mL, clone 6E10, Cat #: 803002, Biolegend).

Tat-tagged peptides were obtained from SYNpeptides, Shanghai, China and consisted of the tat-derived sequence YGRKKRRQRRR, which makes peptides membrane-permeant, and either residues 1923-1942 of Ca_V_1.2 (LGRRASFHLECLKRQKNQGG), which mimics the β_2_AR binding site on Ca_V_1.2 and displaces the β_2_AR from Ca_V_1.2 (termed tat-Pep1923: YGRKKRRQRRRLGRRASFHLECLKRQKNQGG) or the control peptide in which the sequence of residues 1923-1942 had been scrambled (termed tat-Pep1923scr: YGRKKRRQRRRGGQNSFAGLRLERCLHKRQK).

Other chemicals were from the usual sources and of standard biochemical purity. ω-conotoxins GVIA and MVIIC were obtained as synthetic peptides (China Peptides, Wujiang, China; single channel recordings) or from biological sources (gifts from Drs. Samuel Espino and Baldomero Olivera, University of Utah; Ca^2+^ imaging).

### Animals

All procedures followed NIH guidelines and had been approved by the Institutional Animal Care and Use Committee (IACUC) at UC Davis (protocol number 23965) and West Virginia University (#2102040142_R1 and #2105042129). Dissociated hippocampal cultures (HC) were prepared from Sprague-Dawley rat E17-E18 embryos and P0 or P1 pups from S1928A KI mice and WT mice of the same genetic background (50%C57bl/6J and 50%129) (*29, 92*). Rats were obtained from Harlan, Charles River, and Envigo and mice from in house colonies.

### Hippocampal cultures

Hippocampal neurons were cultured as before (*93, 94*) from either wild-type E18 male and female E17-E18 embryos from Sprague Dawley rats or from P0 or P1 WT or S1928A KI mouse embryos. Hippocampi were excised from brains in ice-cold Hank’s Buffer (Sigma-Aldrich, Cat#H2387) with 10 mM HEPES (Gibco Cat #: 15630-080), 0.35 g/L NaHCO_3_ and 5 μg/ml gentamicin (Gibco Cat #: 15710-064) and digested in 0.78 mg/ml papain (Roche, Cat #: 10108014001) in 5 ml of the above Hank’s buffer at 37°C for 2 x 15 min, in an incubator containing 5% CO_2_ and 95% air. The digested tissue was washed two times with culture medium and subsequently triturated in this medium. The culture medium consisted of 1x B27 supplement, 1x Glutamax, 5% FBS and 1 µg/ml gentamicin in Neurobasal medium. Cells were plated at a density of 40,000 – 60,000 neurons/well in 24-well plates on borax coated coverslips and cultured in an incubator at 37°C and 5% CO_2_ and 95% air.

### Cell-attached single channel recordings and analysis

LTCC activity was recorded from HC DIV15-25 in cell-attached mode at an Olympus iX50 inverted microscope as before (*29, 30, 94–96*). The membrane potential was collapsed by using a high K^+^ concentration in the external solution, which contained 120 mM K^+^-Glutamate, 25 mM KCl, 2 mM MgCl_2_, 1 CaCl_2_, 10 EGTA, 10 HEPES, and 2 mM Na^+^-ATP (pH 7.4 KOH). The intracellular pipette solution contained 110 mM BaCl_2_, 20 mM TEA-Cl, and 10 mM HEPES (pH 7.4 with TEA-OH, 325-330 mOsM). 1 µM ω-conotoxins GVIA and 1 µM MVIIC (China Peptides) were included in the pipette to isolate LTCCs by inhibiting N-, P-, and Q-type currents. Recording pipettes were made from borosilicate glass, ID 0.86 mm OD 1.5 mm using a micropipette puller (Sutter Instruments, Model P-97) followed by pipette polishing. Pipette resistance was 7-12 MΩ.

In the initial experiments, transmembrane patch was repeatedly depolarized from a holding potential (HP) of -80 mV to a test potential (TP) to 0 mV for 2 s followed by 5 s HP for ≥ 100 consecutive traces. In later experiments HP was -40 mV to further minimize activity of non-LTCCs, especially R-type channels (*97*). Sampling frequency was 10 kHz. Currents were low-pass filtered at 2 kHz using an Axopatch 200B amplifier with data digitalization using a Digidata 1440A (Axon Instruments) and read into Clampex 10 of pCLAMP 10 software suite (Molecular Devices). Leak subtraction was performed subsequently after data acquisition and event list were created analyzed based on the half-height criterium (*98*). Peak ensemble average currents were created from averaging all sweeps from one neuron and averaging the average traces from all neurons in each experimental group. Ensemble currents depict a temporal superposition of all idealized channel openings of a single experiment at timepoint *t_i_.* The summation of all sampling points I(t_i_) at timepoint *t_i_* over all current sweeps (j) devided by the number of all recorded sweeps (*M_0_*) results in the equation: I*_sum_*(*t_i_*) = ∑_𝑗_ 𝐼 ∗ (𝑡𝑖)𝑗/𝑀0. This summation of all current sweeps allows the visual readout of the local maximum (I_peak_) (*99*).

### Analysis of single channel data

Statistical analyses were performed using GraphPad Prism 10. All data are represented as mean values ± SEM. Data sample sizes, *P* values and statistical tests are indicated in the figure legends. Single channel Po data analysis was performed as previously described (*29, 30, 94–96*). In brief, 500 nM BayK 8644 (Tocris) was used to augment LTCC activity in order to determine the number of LTCCs in each patch (*29, 30, 94–96*). Channel number was determined based on observed simultaneous detected opening events (*k*, stacked current levels) with 500 nM Bay K 8644 present during long consecutive sweep recordings of ≥ 20 min. Recordings with more than 4 channels (k≥4) in the patch were not considered for data analysis due to the likely overinterpretation in Po (*100*). Population distribution and statistical significance was determined based on parametric testing using either a one-way ANOVA with Bonferroni post-hoc correction for more than 2 groups or by an unpaired Student’s T-test. Grubb’s outlier test was used if necessary to exclude outliers (α =0.05). Data sample size was based on a 95% confidence level with 5% accuracy.

Dwell time histograms were calculated and analyzed as previously described (*101*). In brief, the abscissa shows the logarithmic transformation of time, x=lg(t) for the i^th^ time constant, respectively τ1- τ3 for open and τ1- τ4 for closed, to optimally visualize dwell times with their respective fractional proportions (*102*). The ordinate shows the number of detected events given as square root numbers. A Maximum Likelihood Estimate (MLE) optimization algorithm (clampfit 11.3) was used as a best fit model to describe the underlying probability density distribution resulting in individual (dashed) and a sum of exponential fits (full) with minimal error rates. Peaks in the gamma distribution represent transitions between close to open states. Cutoff rate for events was calculated as <166µs at a 2kHz filtering rate. Histograms were binned with 10 bins/decade.

### Ca^2+^ imaging in hippocampal neurons

*DIV 10-18* HC from WT and S1928A KI mice were preincubated with 1 µM ω-CTx GVIA and ω-CTx MVIIC for 1 hour, loaded with 5 μM Cal-520, AM (Ca^2+^ sensor; AAT) for 5 min, and washed with Tyrode buffer (in mM: NaCl 150, KCl 5, HEPES 10, Glucose 10, MgCl_2_ 2, CaCl_2_ 2; pH7.3 with NaOH; 310mOsm) for 10 min. Images were recorded (1 per 5 s) at an inverted microscope iX81 (Olympus) with an iXon Ultra (Andor) camera. HC were depolarized by switching the perfusion to high K^+^ Tyrode (in mM: KCl 90, NaCl 50, HEPES 10, Glucose 10, MgCl_2_ 2, CaCl_2_ 2; pH7.3 with KOH; 310mOsm) for 50 s. After 5 min HC were incubated with 100 nM Aβ for 20 min before a second high K^+^ Tyrode application. Mean fluorescence intensity was calculated by subtracting the background signal from the mean intensity of the region of interest (ROI) with Fiji software (Image J). Relative fluorescence over time was determined by normalizing the fluorescence intensity of each image to the baseline (average of the 5 images prior to depolarization). The area under the curve (AUC) was calculated above the baseline level of the curve (set to 1) using Prism 10 software (GraphPad). Aβ-dependent Ca^2+^ entrance was calculated as AUC_Aβ_/ AUC_control_.

### Detection and analysis of miniature excitatory postsynaptic currents (mEPSC)

HC at DIV 10-20 were treated with Aβ and either vehicle control or inactive Aβ_INV_ before detecting mEPSCs events. External solution contained (in mM): 115 NaCl, 5 KCl, 2.5 CaCl_2_, 1.3 MgSO_4_, 23 dextrose, 26 sucrose, 4.2 HEPES, 0.001 tetrodotoxin and 0.05 picrotoxin (Tocris), pH 7.2, 295-305 mOsm (sucrose) (*103*). Patch pipettes (2– 4 MΩ) were filled with intracellular solution (in mM): 128 K-Gluconate, 10 NaCl, 1 EGTA, 0.132 CaCl_2_, 2 MgSO_4_, 10 HEPES, pH 7.2, adjusted with sucrose to possess 10 mOsm less than the extracellular solution. Data with a series resistance greater than 20 MΩ were discarded. Raw data were collected with Clampex 10.6 by an Axopatch 200B patch-amplifier (Molecular Devices). Neurons were voltage-clamped at -60 mV and whole-cell currents were filtered at 2 kHz and sampled at 20 kHz. The mEPSC events were then analyzed with the template method provided by Pclamp 10.6 and the accuracy of detection was confirmed by visual inspection. Threshold detection level for mEPSC was set larger than 7 pA. Mean values were directly retrieved from the event statistics list. Additionally, cumulative plots were depicted with 5 bins/decade (amplitude) and 0.2 bins/decade (inter-event interval).

### Slice preparation and treatment, immunoprecipitation (IP) and immunoblotting

6 to 12-week-old mice were decapitated, and brains placed into ice-cold slicing buffer (in mM: 10 NaCl, 230 sucrose, 26 NaHCO_2_, 1.2 KH2PO_4_, 2.5 KCl, 1 CaCl_2_, 1.3 MgSO_4,_ and 10 D-glucose, 290-300mOsm/kg, saturated with 95% O_2_ and 5% CO_2_; final pH 7.3). About one third of the front and back of the brain were removed and 400 µm thick slices containing hippocampus prepared with a vibratome (Leica VT 1000A) and equilibrated in artificial Cerebrospinal Fluid (ACSF; in mM: 126 NaCl, 26 NaHCO_2_, 1.2 KH2PO_4_, 2.5 KCl, 1 CaCl_2_, 1.3 MgSO_4_ and 10 D-glucose, 290-300mOsm/kg, saturated with 95% O_2_ and 5% CO_2_; final pH 7.3) for 1.5 hrs. at 32°C before use.

Slices were then treated with 1 μM Aβ or Aβ_INV_ plus 100 nM ICI118551 (preincubate for 5 minutes) if indicated for 25 min before trituration with insulin syringes in 250 μl solubilization buffer (1% Triton X-100, 150 mM NaCl, 10 mM EDTA, 5 mM EGTA, 50 mM Tris-HCl, pH 7.4) (*26*) containing protease inhibitors (10 μg/ml pepstatin A, 1 μg/ml leupeptin, 2 μg/ml aprotinin and 200 nM PMSF; PMSF was added right before trituration and again at the start of IP), and phosphatase inhibitors (4 μM microcystin LR, 1 mM *p*-nitrophenyl phosphate, 25 mM Na-pyrophosphate, and 25 mM NaF) (*104, 105*). Insoluble material was removed by ultracentrifugation (40,000 rpm for 30 min) followed by IP (2.5 hrs on ice) with our FP1 antibody against Ca_V_1.2 using protein A Agarose. The beads would then be washed first by a low salt washing buffer (0.05% Triton X-100, 150mM NaCl, 10mM EDTA, 10mM EDTA, 10mM EGTA, and 10mM Tris-HCl (PH 7.4)), then by a high salt washing buffer (750mM NaCl, 10mM EDTA, 10mM EDTA, 10mM EGTA, and 10mM Tris-HCl (PH 7.4)), and finally with a no-salt washing buffer (10mM EDTA, 10mM EDTA, 10mM EGTA, and 10mM Tris-HCl (PH 7.4)). Samples underwent SDS-PAGE and were transferred onto PVDF membranes. Immunoblots (IBs) were blocked with 5% or 10% dry milk powder (depends on the type of the antibodies) in TBS plus 0.1% Tween-20 (TBST), incubated with primary antibodies (5% or 10% dry milk in TBST; 4 °C overnight), washed three times with TBST, incubated with HRP-conjugated secondary antibody (1:10,000 in TBST; room temperature for 1hr), and washed several times for 45 min before detection of HRP signals with ECL or ECL plus chemiluminescence reagents by film exposure. Multiple exposures of increasing length ensured that signals were in the linear range, as described (*18, 104*).

### Surface biotinylation

Surface biotinylation experiments were performed and analyzed essentially as before(*96, 106*). Slices were treated in ACSF with 1 μM Aβ or Aβ_INV_ for 25 min before being washed twice with ACSF to remove the drugs. They were then incubated with 1 mg/ml Sulfo-NHS-SS-biotin (Pierce) dissolved in ACSF for 30 min on ice. Sulfo-NHS-SS-biotin was quenched by washing slices two times with ACSF and then one time with ACSF containing 100 mM glycine dissolved in filtered PBS. After the quenching, slices were washed two more times with ACSF. Slices were extracted and samples centrifuged as above and the cleared supernatant then incubated with NeutrAvidine (NtrAv) Sepharose beads overnight at 4°C. The NtrAv beads were pelleted and washed three times with solubilization buffer followed once with PBS. Bead-bound proteins were extracted with SDS-PAGE sample buffer and fractionated by SDS-PAGE (7.5 % precast acrylamide gradient gels, BioRad), transferred to membranes and subjected IB as described above. Assessment and quantification was done as described (*96*).

### Analysis of LTP in Hippocampal slices

Recordings were as detailed earlier (*30, 107*). Briefly, 400 µm thick coronal forebrain slices were prepared from 6-16 months old WT and S1928A KI mice in ice-cold buffer containing (in mM): 2.5 KCl, 3 MgSO_4_, 1.2 NaH_2_PO_4_, 26 NaHCO_3_, 10 dextrose, 220 sucrose, 1.1 CaCl_2_, follow by incubation for at least 1 h at 32°C before placing into the recording chamber. Slices were perfused with a flow rate of 1 ml/min at 30°C with artificial cerebrospinal fluid (ACSF; in mM: 127 NaCl, 1.9 KCl, 1 MgSO_4_, 1.2 NaH_2_PO_4_, 26 NaHCO_3_, 10 dextrose, 2.2 CaCl_2_) equilibrated with 95% O_2_ and 5% CO_2_ (final pH 7.3) with a peristaltic pump but without recirculation. Schaffer collaterals in CA1 were stimulated in stratum radiatum near recording electrode every 15 s with a tungsten concentric bipolar microelectrode and fEPSPs recorded with a glass electrode filled with ACSF. Signals were amplified with an Axopatch 1D (Axon Instruments), digitized (Digidata 1320A, Axon Instruments), and analyzed (Clampex 9, Molecular Devices). Stimulus strength was set to about ∼50% of the maximal response as determined in input-output response curves (typically 0.2-1.0 mV). Baseline recordings were performed until fEPSP responses were stable. Aβ or Aβ_INV_ were applied at 1 μM for 20 min or 200 nM for 1 h alone or together with indicated drugs. Then, LTP was induced by two 1 s-long tetani of 100 Hz that were 10 s apart and fEPSPs recorded for another 50-60 min. The means of the initial slopes of fEPSPs during the 10 min preceding the tetani corresponded to 100% baseline level and the means of fEPSP initial slopes obtained 60 min after the tetani determined potentiation relative to the 100% baseline.

### Cortical cultures and transfection

Dissociated cortical neurons were prepared from the embryonic day 17-18 (E17-18) male and female rat cerebral cortices and cultured for 18-25 days in vitro (DIV) in Neurobasal medium with phenol red (cat#: 12348017, Thermo Fisher Scientific) supplemented with 1x B27 supplement (cat #: 17504044, Thermo Fisher Scientific), 200 mM L-glutamine (cat #: 25030081, Thermo Fisher Scientific) and 0.1 mg/mL penicillin-streptomycin (cat #: 15140122, Thermo Fisher Scientific) as described previously (*53, 70*). Neurons were plated on poly-D-lysine (cat #: 354210, Corning, Corning, NY) and laminin (mouse; cat #: 354232, Corning) coated glass coverslips (12 mm, #1.5; cat #: 64-0732, Warner Instruments, Camden, CT). Neurons were plated at 150,000 cells/well in 24-well plates and were maintained in a humidified 37°C incubator with 5% CO_2_.

Cortical neurons were transfected at DIV 3 as previously described (*53, 70*) using Lipofectamine 2000 (cat#: 11668027, Thermo Fisher Scientific). EGFP, under the control of a human ubiquitin promoter (pFUg-EGFP), was used as a cell filling dye to visualize neuronal morphology (*53, 70*). Briefly, for each well of a 24-well plate, the conditioned medium was first collected from plated neurons and replaced with 300 µl of Neurobasal medium without any supplements. 100 µl of transfection mix containing 0.5 µl of Lipofectamine 2000 and 200 ng of pFUg-EGFP plasmid was then added to each well of a 24-well plate. Neurons were incubated with the transfection cocktail at 37°C for 2 hours. After 2 hours, the transfection medium was replaced with 500 µl of the warmed conditioned medium that was sterilized by passing through a 0.2 µm filter (Millipore-Sigma). Transfected neurons were then placed in a humidified 37°C incubator until DIV 21-25, at which point they were treated with Aβ and processed for immunocytochemistry and STED imaging.

### Immunocytochemical analysis of cortical cultures

Cortical cultures were pretreated at 18-21 DIV with vehicle, ICI 118551 (400 nM, Cat#: 0821, Tocris), or isradipine (10 µM, Cat#: 2004, Tocris) for 1 h or tat-Pep1923 or tat-Pep1923scr (both at 1 µM) for 30 min to 20-24 DIV cortical cultures before addition of 500 nM Aβ (ICI and isradipine) or 100 nM Aβ (tat peptides) for a 24 h incubation period at 37°C in a humidified cell culture incubator supplied with 5% CO₂. Cultures were fixed in 4% PFA containing 2% sucrose supplemented in 1xPBS for 8 min at room temperature, washed three times with 1xPBS, blocked and permeabilized for 1 h at room temperature in 1% ovalbumin (cat#: A5503, Millipore-Sigma) and 0.2% gelatin from cold-water fish (cat#: G7041, Millipore-Sigma) in 1xPBS containing 0.1% saponin (cat#: 558255, Millipore-Sigma) (*53, 70*). Neurons were immunostained for 2 hours at room temperature or overnight at 4°C with chicken anti-GFP (1:2000, cat#: ab13970, Abcam, RRID# 300798) to visualized neuronal morphology and dendritic spines, FluoTag-X2 anti-PSD-95 (nanobody) directly conjugated to Abberior STAR 635P (1:500, cat#: N3702, NanoTag Biotechnologies, RRID#: 3076102) and mouse IgG2b anti-Bassoon (1:500, cat#: 75-491, Neuromab, RRID#: 2716712) to visualize synaptic nano-organization. Neurons were then washed three times with 1xPBS, and exposed to corresponding secondary antibodies (Goat anti-chicken-Alexa Fluor 488, cat#: 103-545-155, Jackson Immunoresearch, RRID: 2337390; Goat anti-mouse IgG2b-Alexa Fluor 594, cat#: 115-585-206, Jackson Immunoresearch, RRID#: 2338886) diluted 1:500 in the blocking buffer for 1 h at room temperature. After washing three times with 1xPBS, coverslips were mounted with ProLong Glass antifade mounting medium (cat#: P36984, Thermo Fisher,) before confocal and STED imaging after 24-48 hours, when the mounting medium was fully cured.

### Surface staining of AMPARs

Cortical neurons (DIV 18-21) were pre-treated for 30 min with vehicle, 400 nM ICI118551, or 10 µM isradipine before addition of either 100 nM Aβ or vehicle for 20 min, and live surface labeling with antibodies for GluA2 (Fab-151F) directly conjugated to Atto647N and rabbit anti-GluA1 in in Neurobasal medium containing B27 and glutamine for 20 minutes at 37°C (*50*) (both gifts from Dr. Daniel Choquet, Univetsity of Bordeaux, France). Neurons were washed three times with warmed 1xPBS, fixed for 8 min, and blocked for 1 h as above. Neurons were then immunostained either for 2 h at room temperature or overnight at 4°C with chicken anti-GFP (1:2000, cat# ab13970, Abcam, RRID# 300798) and anti-PSD-95 antibody (IgG1 clone K28/43, cat# 75-02, Neuromab). After three additional washes with 1x PBS, coverslips were incubated for 1 hour at room temperature with anti-rabbit conjugated to Alexa Fluor 594 (cat# 111-585-144, Jackson Immunoresearch), anti-mouse (IgG1) conjugated to Alexa Fluor 555 (cat# A21127, Thermo) and anti-chicken conjugated to Alexa Fluor 488 secondary antibodies diluted 1:500 in the blocking buffer to visualize GluA1, PSD-95 and GFP, respectively. After final three washes in 1x PBS coverslips were mounted on slides using ProLong Glass antifade mounting medium (Thermo) to be used for STED imaging.

### STED imaging

Two-color tau-STED imaging of endogenous proteins was performed on fixed neurons in cell culture using protocols developed previously (*53, 70, 71*). A Leica Stellaris 8 3D tau-STED confocal and super-resolution system (Leica Microsystem, Mannheim, Germany) equipped with a tunable white light laser (WLL), CW 592 nm, CW 660 nm and pulsed 775 nm STED depletion lines was used for image acquisition. HyD-X detectors set to photon counting mode at a 12 mV threshold were used to capture single photons for STED image acquisition. The 100x oil immersion objective (1.4 NA) with 4.5x digital zoom to obtain ∼20 nm pixel size was used to acquire image stacks at 150 nm intervals using 400 Hz scanning of 1024 x 1024-pixel imaging fields (∼ 20 µm x 20 µm). Line accumulations (2x – 3x) were used to image individual channels. Stacks for each channel were acquired in a sequential mode using the “between stacks acquisition” setting. Target proteins (PSD-95 and Bassoon) labeled with Abberior STAR 635P and Alexa Fluor 594 were acquired using the Fast Lifetime Contrast (FALCON) enabled tau-STED module (Leica) with the time-gate on HyD-X detectors adjusted between 0.1 and 6 nanoseconds. Abberior STAR 635P labeled endogenous proteins were excited with the 635 nm laser (10-15% maximal AOBS laser power). Alexa Fluor 594 labeled endogenous proteins were excited using the 594 nm laser (5-12% maximal AOBS power). The pulsed 775 nm STED depletion line set at 10% of maximal AOBS laser power was used for STAR 635P and 15% of maximal AOBS laser power for Alexa Fluor 594 to generate STED. All data shown were imaged using 3D STED with 20% STED laser power re-directed toward the Z-doughnut. For simultaneous confocal/STED imaging, fluorophores that were not exposed to STED lasers (Alexa Fluor 488) were imaged first, using only excitation lines provided by the WLL. This procedure was then followed by sequential STED acquisition of Abberior STAR 635P and Alexa Fluor 594 fluorophores.

### Image processing

Following image acquisition, background photons were removed from raw STED images by adjusting tau strength using the tau-STED module in the FALCON application on the Stellaris 8 STED instrument (Leica Microsystems) (*71*). The ability to remove unwanted photons relies on generating fluorescence lifetime imaging (FLIM) profiles of every photon captured by HyD-X detectors (*71*). The tau strength was adjusted to reduce blur surrounding cluster edges in a manner that did not distort existing clusters or create cluster artifacts. With 5-10 captured photons per pixel, this usually resulted in a tau strength between 100-200. Once the tau strength was determined for each STED channel, it was kept constant throughout all experiments. Afterward, tau-STED images were subjected to 0.8-pixel Gaussian blur, and brightness was adjusted to generate high-contrast images for figures and analysis. None of the images acquired were deconvolved. Tau-STED images were obtained as 16-bit Tiff files. These images were initially adjusted in ImageJ by subtracting the background (mean pixel intensities of the entire 1024 x 1024 frame) from every channel. We then converted these 16-bit files to 8-bit images that were subsequently used in analysis.

### STED analysis

All analyses were conducted offline using Fiji Image J and built-in macros on a per spine basis as described below and previously published (*53, 70, 71*). Spines for analysis were collected from independent dendritic segments (25-50 µm) in a minimum of three GFP transfected neurons. Briefly, images of GFP labeled spines were acquired at confocal resolution (∼250 nm), and a 0.8-pixel Gaussian blur was applied to filter out noise. Individual spines were converted to binary masks by thresholding GFP (enhanced by chicken GFP immunolabeling) to the mean +2x SD of the 1024 x 1024-pixel area corresponding to the entire image field. The localization of PSD-95 and Bassoon nanoclusters was then determined per spine. Nanoclusters of synaptic proteins (acquired in tau-STED super-resolution) were identified by binarizing each channel separately using intensity thresholds. Thresholds were defined as the mean +2x SD of intensity values of a 50 x 50 image pixel area that typically corresponded to the area of spine head. In each spine ROI, STED-resolved nanoclusters were defined as a minimum of 10 and a maximum of 100 continuous pixels corresponding to an area of 0.002 – 0.15 µm^2^. The separation between neighboring STED resolved clusters was identified from the line intensity profiles of nearby clusters and was defined as the mean +1.5x SD of a local 50 x 50 pixel area that approximately corresponded to the maximum size of a spine head. The resulting thresholded nanomodules were used to determine whether these modules colocalized with individual spines. All spines on a single dendritic segment in the image were subjected for analysis.

### Analysis of synaptic AMPAR nanoclusters

Three-color tau-STED images of GluA1, GluA2, and PSD-95 were analyzed as maximum intensity projections in an unbiased manner with a custom-built ImageJ macro. Briefly, the brightness of each maximum projection image was adjusted individually by subtracting the background, defined as the average intensity of all pixels in the image. The maximum intensity of each image was then normalized to 50% of the highest pixel intensity value. Following this adjustment, 16-bit tau-STED images were converted to 8-bit images and subjected to a one-pixel Gaussian blur. First, GluA1, GluA2, and PSD-95 were thresholded to identify 90–95% of all clusters in the images. All thresholded images were then analyzed for particles containing between 10 and 500 contiguous pixels and watershed was applied to separate merged clusters. Next, we determined the colocalization between GluA1 and GluA2 nanoclusters. To identify GluA1-lacking and GluA2-lacking nanoclusters, we subtracted the colocalized GluA1/GluA2 masks from the original GluA1 and GluA2 tau-STED images, respectively. These subtracted images were then thresholded to the same values as those used for the colocalized masks. Finally, the colocalized GluA1/GluA2, GluA1-lacking, and GluA2-lacking masks were subjected to colocalization analysis with PSD-95 nanocluster masks to quantify synaptic AMPARs for statistical analysis.

### Cytotoxicity test of HCs

Cytotoxicity assays were performed essentially as described (*108*). HCs were treated for 48 h with vehicle or 1 μM Aβ and various drugs, incubated with membrane-impermeant propidium iodide (10 μg/ml; Sigma-Aldrich) and membrane-permeant Hoechst 33342 (10 μg/ml; Molecular Probes, Eugene, OR), washed twice with PBS, fixed with 4% paraformaldehyde in PBS, mounted, and imaged on a BX50 epifluorescence microscope (Olympus).

## ACKNOWLEDGEMENTS

We thank Drs. Baldomero Olivera and Samuel Espino (University of Utah) for the ω-conotoxins GVIA and MVIIC used in the Ca^2+^ imaging experiments; Dr. Daniel Choquet (University of Bordeaux, France) for the GluA1 antibody and the GluA2 Fab-151F - Atto647N probe that were used for surface staining; and Drs. Kenneth Ginsburg and Timothy Hanks, UC Davis, and Dr. Andrew Plested (Humbold University, Berlin, Germany) for feedback on data analysis.

This work was supported by NIH grants RF1 AG055357 (JWH), R01 NS123050 (JWH), and P20 GM109098 (MH), the Alzheimer’s Association grant AARG-NTF-23-1150820 (MH) and NSF grant OIA-2242771 (MH) with additional support by NIH grants T32 GM007377 and F99 NS135807 (ZME-T), T32 MH082174 and T32 GM099608 (AAJ), T32 GM144303 (RAB), R01 GM129376, R01 MH134119, and VA merit award IK6BX005753 (YKX), R01 HL121059 and R01 HL149127 (MFN).

## Author contributions

PB: conceptualization, data curation, formal analysis, investigation, methodology, project administration, data evaluation, visualization, writing - original draft preparation, writing and editing.

SR: investigation, methodology, formal analysis.

JDS: investigation, methodology, formal analysis.

ZZ: methodology, formal analysis.

ZME: methodology.

JP: investigation, methodology, formal analysis.

AAJ: methodology.

RAB: methodology.

SYH: methodology.

AA: methodology.

YKX: review, discussion.

CYC: supervision, methodology, review, discussion

MNC: writing - review.

MFN: supervision, review.

MCH: supervision, methodology, writing - review and editing.

MH: investigation, methodology.

JWH: conceptualization, funding acquisition, investigation, methodology, project administration, supervision, writing - original draft preparation, writing - review and editing.

## Data and materials availability

All data needed to evaluate the conclusions in the paper are present in the paper or the Supplementary Materials.

---

**fig. S1.**
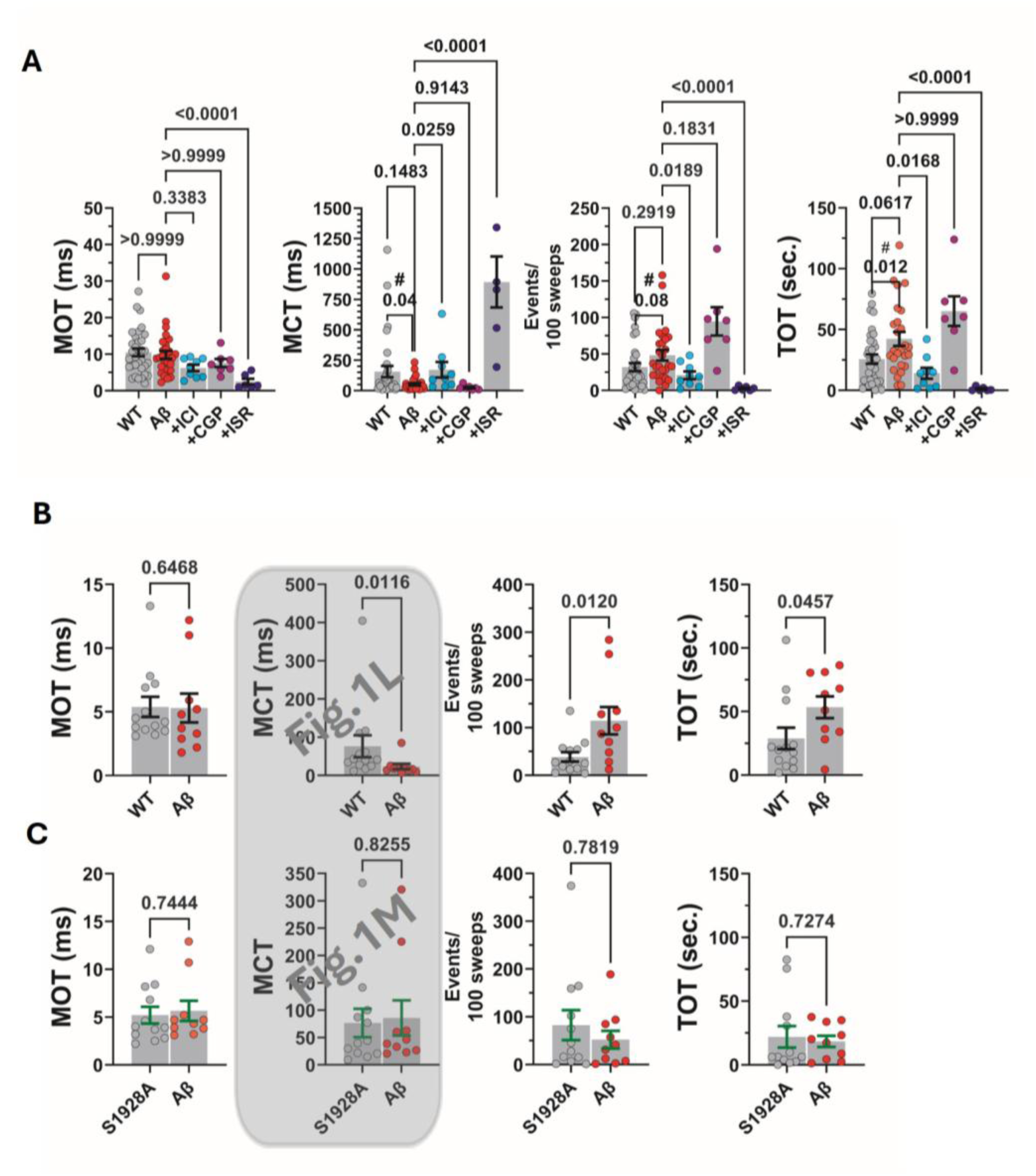
Acute effects of Aβ on overall Ca_V_1.2 single channel activity parameters. A-C : Quantification of effect of Aβ on population data for mean open time (MOT), mean closed time (MCT), event frequency, and total open time (TOT) for recordings from rat HCs (**A**). The increase in Po in rat HC (Fig. 1) can mainly be attributed to a reduction in channel MCT whereas MOT is unaffected. The reduction in mean closed time means that the channel gates at a higher channel opening frequency, overall resulting in a longer TOT. The same is true for WT mouse HC neurons (**B**) whereas none of these parameters are affected in S1928A neurons (**C**), which do not show a change in Po (Fig. 1K).

**fig. S2.**
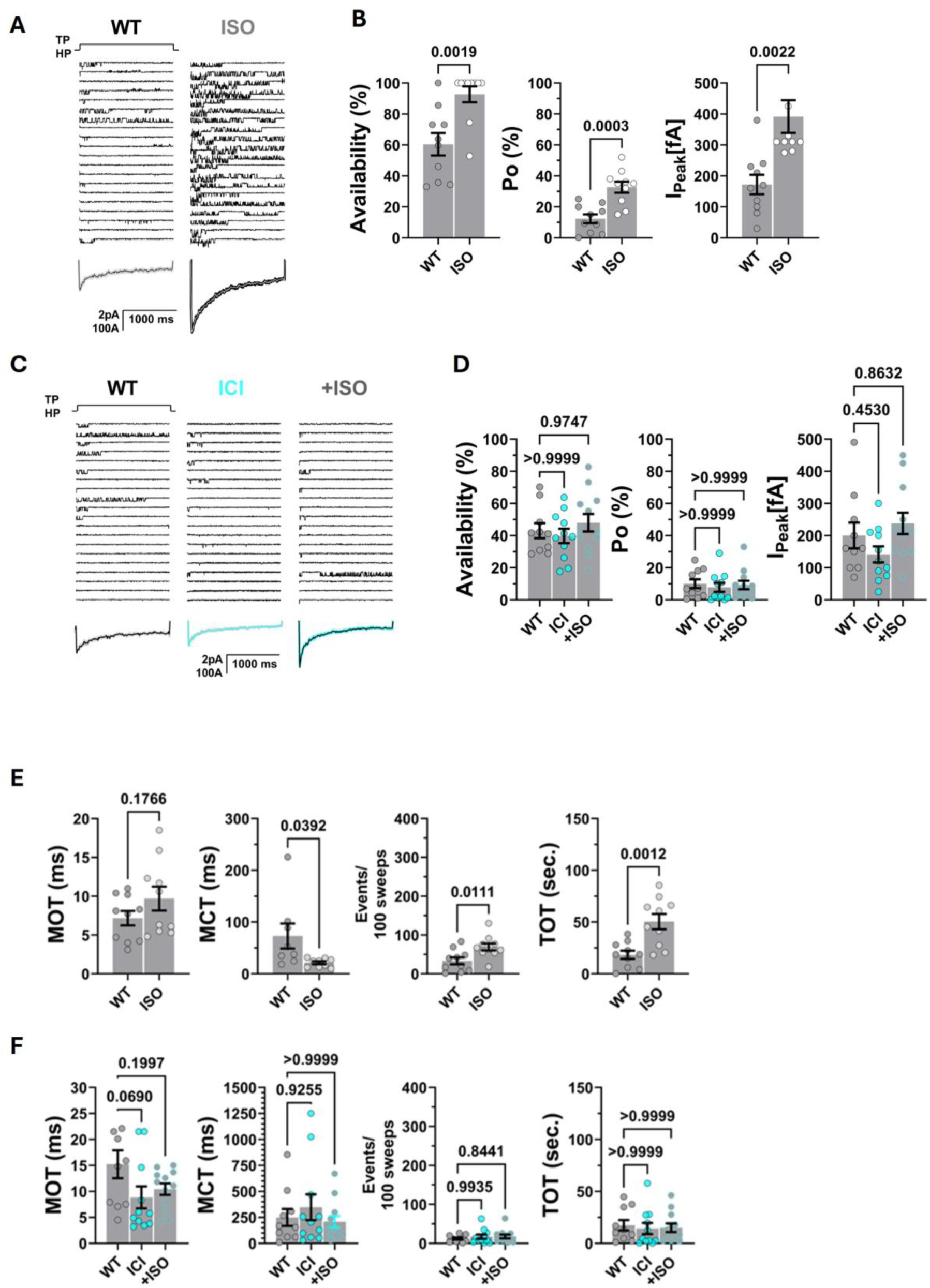
Effects of isoproterenol on overall Ca_V_1.2 single channel activity parameters. To compare theAβ effects on Ca_V_1.2 channel properties with those seen for established βAR agonists, the effects of the βAR agonist isoproterenol (ISO; 1µM) and β_2_AR inverse agonist ICI118551 (ICI; 100 nM) on the different channel paramters were extracted from previously published singe channel recordings (*30*) . Recordings had been obtained from 15-20 DIV mouse HCs (*30*) (**A,B**) and rat HCs (**C,D**) upon depolarizations from a holding potential (HP) of -80 mV to the a test potential (TP) of 0 mV for 2 s. **A**: Twenty consecutive exemplary traces. The pipette solution contained either internal solution alone (WT) or 1µM ISO. **B:** Quantification of population data after correcting for the number of observed simultaneous openings (k≤4). Parameters of availability, Po, and I_peak_ with and without ISO. **C:** Twenty consecutive exemplary traces for untreated rat neurons, 100nM ICI118551 or 100nM ICI plus 1 µM Iso. **D:** Quantification of population data after correcting for the number of observed simultaneous openings (k≤4). ICI alone showed no effect on baseline LTCC avtivity. **E, F:** Quantification of MOT, MCT, TOT, and opening frequency. Significance was tested with an unpaired T-test (**B, E**) or an ANOVA with Bonferroni correction (**D, F**), p<0.05. All data are given as mean ± SEM.

**fig. S3.**
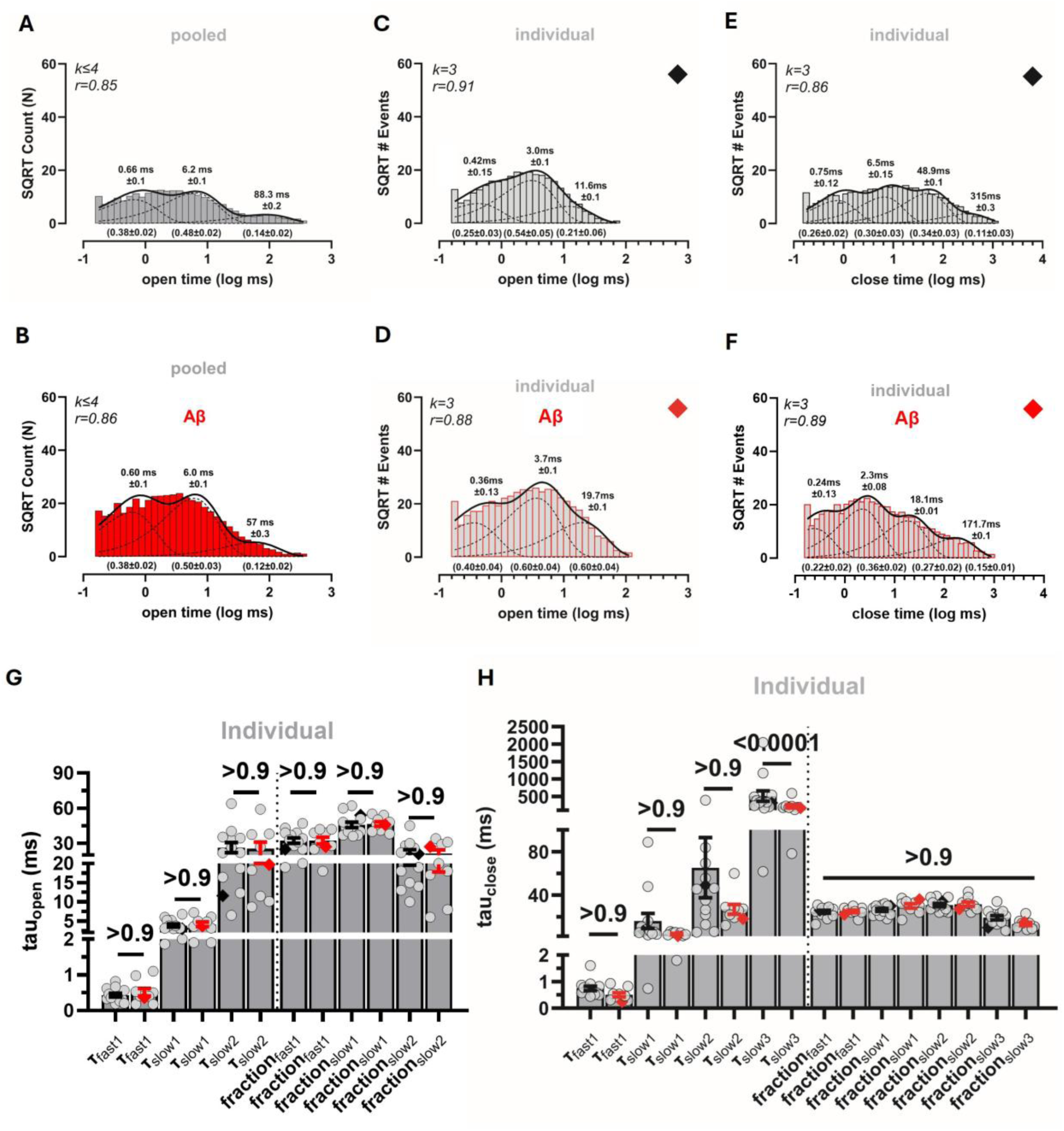
Effects of Aβ on open and closed state dwell time distributions for Ca_V_1.2 single channel activity. **A-F**: Log-binned dwell time histograms for LTCC single channel open (**A-D**) and closed time (**E, F**) interval distribution for recordings from WT neurons for control (**A, C, E**) and 10 µM Aβ (**B, D, F**; for pooled closed time analysis see Fig. 1N). Histograms depict open and close distributions for all events after data had been pooled for all experiments before data fitting to define different states (pooled; **A, B**) or for an individual experiment with k=3 active channels (**C-F**). The filter rise time (*t_r_*) of our Filtering system (3dB, 4-pole Bessel) was calculated to be ∼166µs, based on *t_r_=0.3321/ƒ_c_* at a 2kHz cut-off frequency (*ƒ_c_*). The correlation coefficient *r* is >0.85 for all analyses. **G, H**: Population data of channel dwell times (**left in each bar diagram**), and their fractional distribution (**right**), which were determined for each individual experiment and are represented by the individual dots. Black diamonds depict experiments shown in C and E under untreated conditions and red diamonds in D and F under treated conditions. All data are means ± SEM (p was determined by One-Way ANOVA and Bonferroni correction, p<0.05). In all individual recordings and also independently analyzed pooled recordings, distributions of open times were best fitted with three exponentials and of closed times with four exponentials. Values are comparable for analysis of pooled and individual data. Aβ did not significantly change open time distribution but reduced closed time constants especially for the slowest tau values for WT neurons (see also Fig. 1N for pooled closed time distribution).

**fig. S4.**
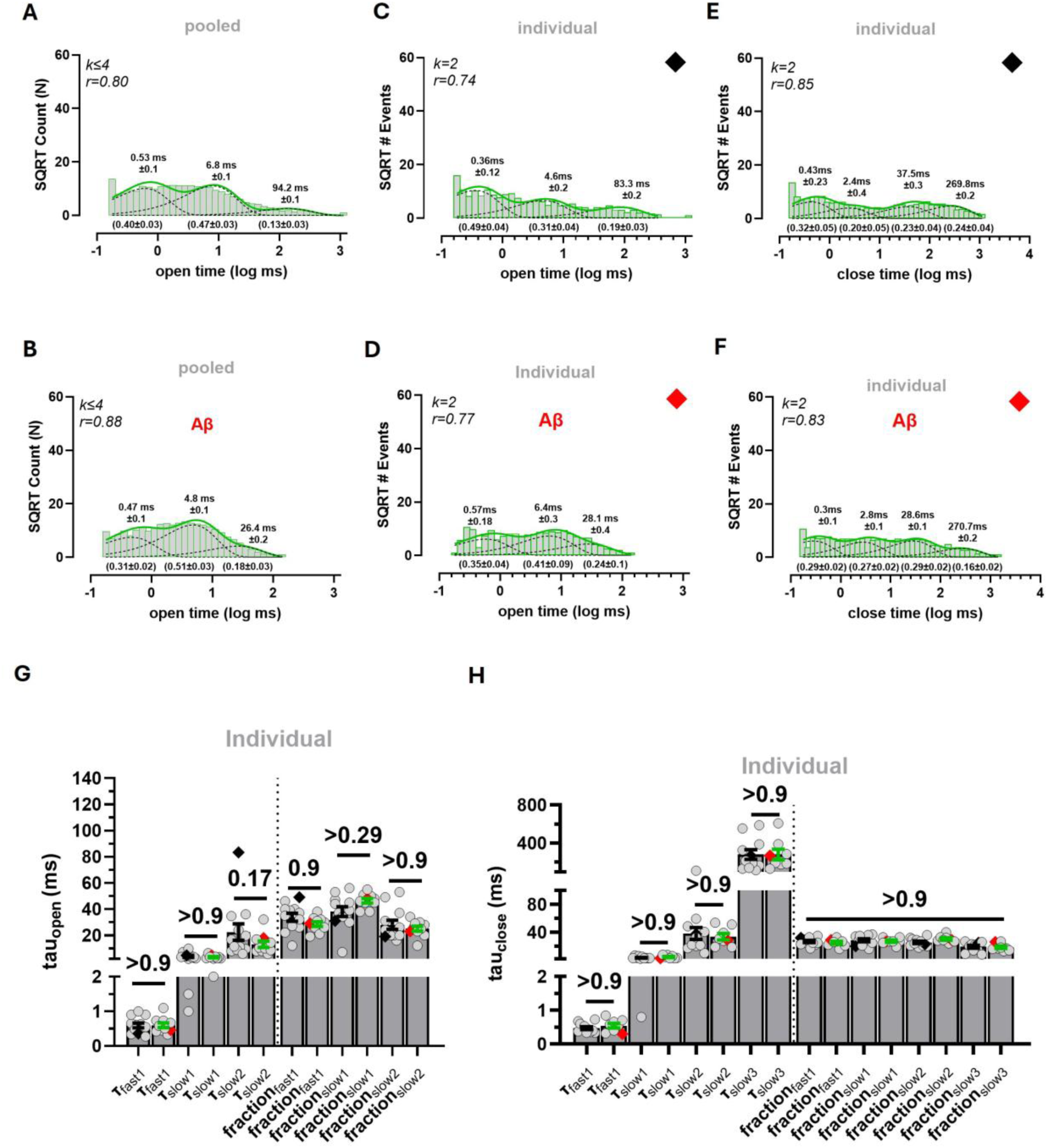
Effects of Aβ on open and closed state dwell time distributions for S1928A mutant Ca_V_1.2 single channel activity. **A-F**: Log-binned dwell time histograms for LTCC single channel open (**A-D**) and closed time (**E, F**) interval distribution for recordings from S1928A KI neurons for control (**A, C, E**) and 10 µM Aβ (**B, D, F;** for pooled closed time analysis see Fig. 1K). Graphs depict distribution for all events after data had been pooled for all experiments before data fitting to define different states (pooled; **A, B**) or for an individual experiment with k=2 active channels (**C-F**). The correlation coefficient *r* is ≥0.74 for all analyses. **G, H**: Population data of channel dwell times (**left in each bar diagram**), and their fractional distribution (**right**), which were determined for each individual experiment and are represented by the individual dots. Black diamonds depict experiments shown in C and E under untreated conditions and red diamonds in D and F for treated conditions. All data are means ± SEM (p was determined by One-Way ANOVA and Bonferroni correction). In all individual recordings and also independently analyzed pooled recordings, distributions of open time were best fitted with three exponentials and of closed times with four exponentials. All values are comparable for analysis of pooled and individual data. Aβ did not significantly change either open or closed time distribution in S1928A KI neurons (see also Fig. 1O for pooled closed time distribution).

**fig. S5.**
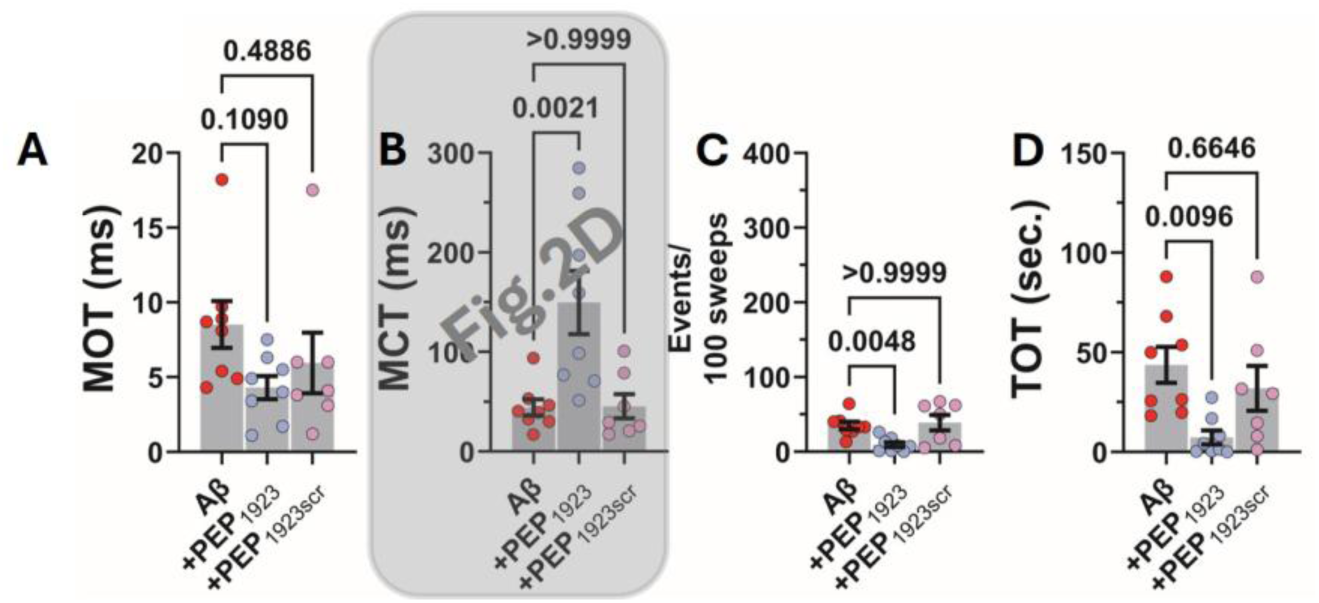
Displacement of β_2_AR from Ca_V_1.2 rescued Aβ-induced changes in single channel parameters. A-D : Quantification of effect of tat-Pep1923 and tat-Pep1923_SCR_ on population data for mean open time (MOT; **A**), mean closed time (MCT; **B**; copied from Fig. 2 for comparison), event frequency (**C**), and total open time (TOT; **D**).

**fig. S6.**
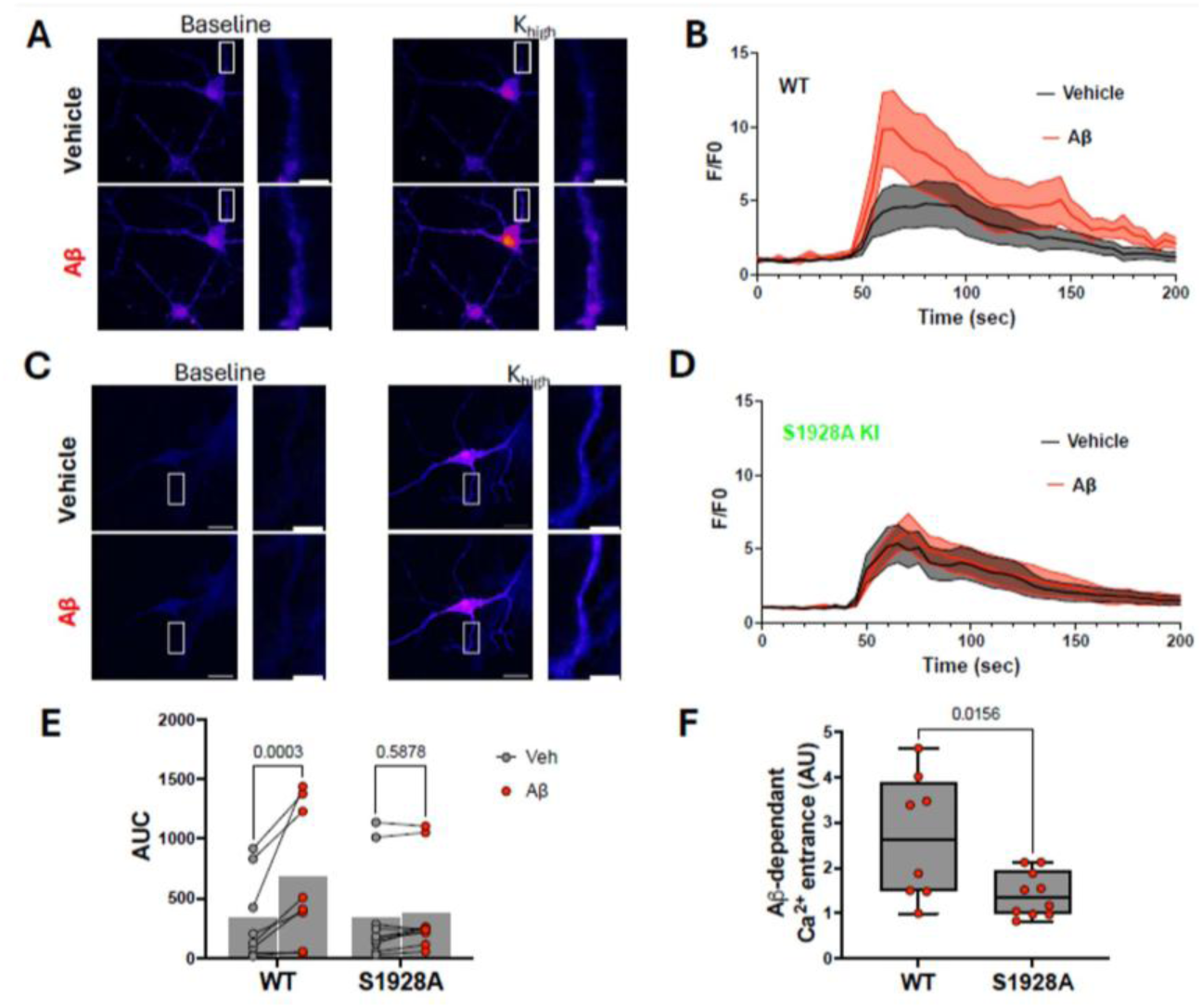
Augmentation of Ca^2+^ influx into dendrites through Ca_V_1.2 by Aβ required phosphorylation of Ca_V_1.2 on S1928. **A-D**: HCs from WT (**A,B**) and S1928A KI mice (**C,D**) at 14-18 DIV were preincubated with 1 µM of the N-, P-, and Q-type channel blockers ω-CTx GVIA and ω-CTx MVIIC and loaded with 5 μM AM-Cal-520 for imaging of Ca^2+^ influx induced by bath application of 90 mM K^+^ before and after application of Aβ. **Left panels** in **A and C** show images before and **right panels** after application of 90 mM K^+^ under the indicated condition on left of each row. Left images in **A and C** show low magnification of neurons (scale bar: 20 µm) and right images enlargements of the labeled fields (scale bar: 5 µm). **B and D** show amalgamated data of all time courses of changes in fluorescence for WT and S1928A KI neurons upon application of 90 mM K^+^ at time point 0. It took about 45 s before the perfusion solution reached the neurons. **E**: Areas under the curves for all recordings (AUC). Bars represent means ± SEM (n: number of neurons recorded from a minimum of N=3 independent experiments; ***p≤0.001, paired t-test). Application of 100 nM Aβ significantly augmented Ca^2+^ transients in WT but not S1928A KI neurons. **F**: Box plots of differences between control and Aβ treatment for each recording from WT and S1928A KI mice (*p≤0.05, unpaired t-test).

**fig. S7.**
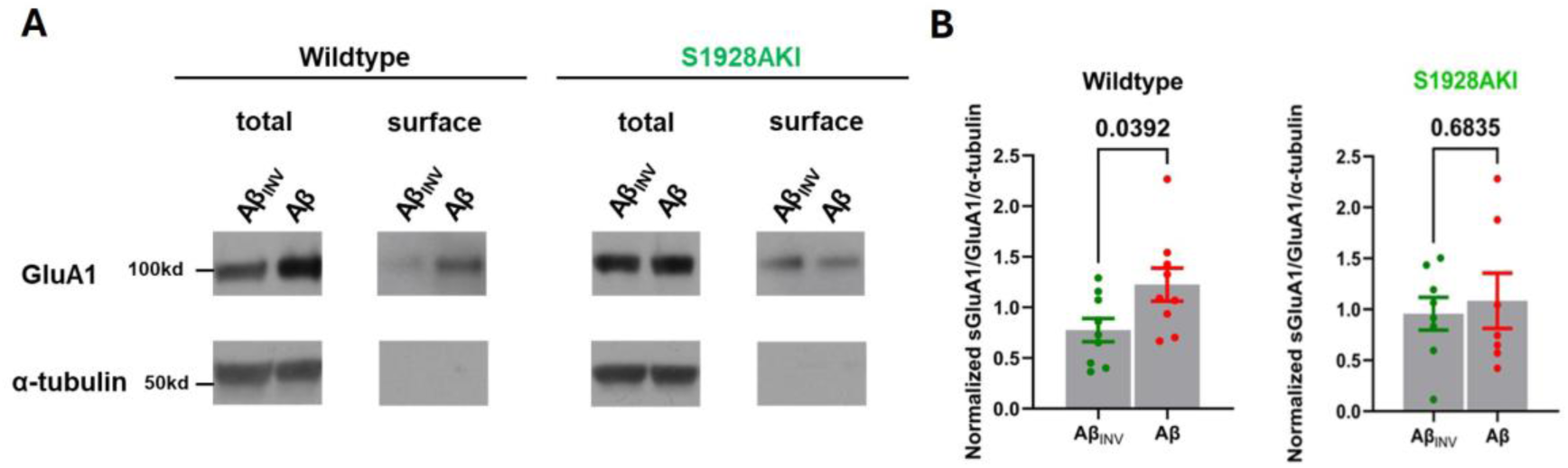
Stimulated of GluA1 trafficking to the surface by Aβ required phosphorylation of Ca_V_1.2 on S1928. Forebrain slices from WT and S1928A KI mice were treated for 25 min with 1 µM Aβ or control Aβ_INV_ before surface biotinylation of Ca_V_1.2 and immunoblotting. **A**: IB for total GluA1 in lysate (**left**) and surface GluA1 after biotinylation and NeutrAvidin Sepharose pull down (**right**). Probing for tubulin served as a loading control to correct for any variability in amounts of protein in lysate samples as well as membrane leakage of the biotinylating reagent. Absence of tubulin in the neutravidin bead pull-down indicated that intracellular proteins had not been biotinylated. **B**: For quantification of immunosignals, first, GluA1 signals in total lysate were divided by tubulin signals to correct for variability between samples. Then, signals for surface GluA1 were divided by the corrected total GluA1 signals to correct for any changes in overall GluA1 content. Each dot in the bar diagram represents this value for one slice from 7 different WT and 4 different S1928A KI mice. Aβ increased surface GluA1 in WT but not S1928A KI mice. All data: mean ± SEM (p values from unpaired t tests).

**fig. S8:**
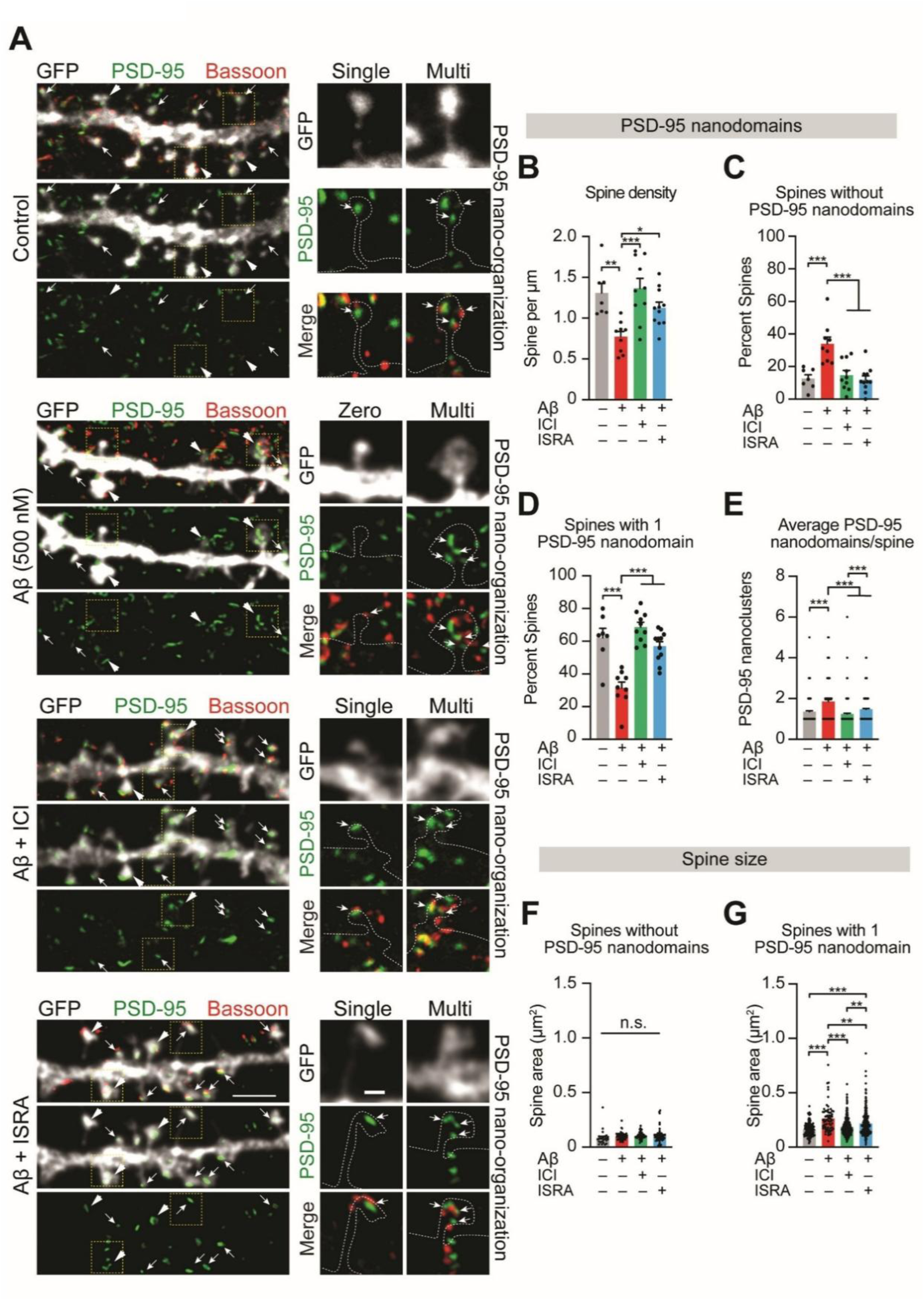
Inhibition of β_2_ARs and LTCCs rescued Aβ-induced changes in spine nanoarchitecture. **A**: Representative tau-STED images of spines from DIV 21–25 cortical neurons transfected with GFP white) and labeled for PSD-95 (green) and Bassoon (red) after treatment with vehicle or 500 nM Aβ for 24 h ± ICI118551 (400 nM) or Isradipine (10 µM). Arrows indicate spines with one aligned PSD-95/Bassoon nanodomain and arrowheads spines with multiple nanodomains. Enlarged views of selected spines (yellow squares) with indicated PSD-95 nanodomain numbers are shown to the right. **B-E**: Overall spine density (**B**), and percentage of spines with 0, 1, or ≥2 PSD-95 nanodomains. Data points represent individual neurons (n = 7 neurons for vehicle, 9 for Aβ, 10 for Aβ+ICI, 11 for Aβ+ISRA). (**F, G**) Average size of spines with 0 or 1 PSD-95 nanodomain. Dots on graph represent individual spines (n for 0 nanodomains for vehicle: 144, Aβ: 67, Aβ+ICI: 263, Aβ+ISRA: 227; n for 1 nanodomain for vehicle: 53, Aβ: 70, Aβ+ICI: 65, for Aβ+ISRA: 124). All data: mean ± SEM from at least 7 different neurons from 3 independent experiments (*p<0.05, **p<0.01, ***p<0.0001, ANOVA, Tukey’s post-hoc).

**fig. S9:**
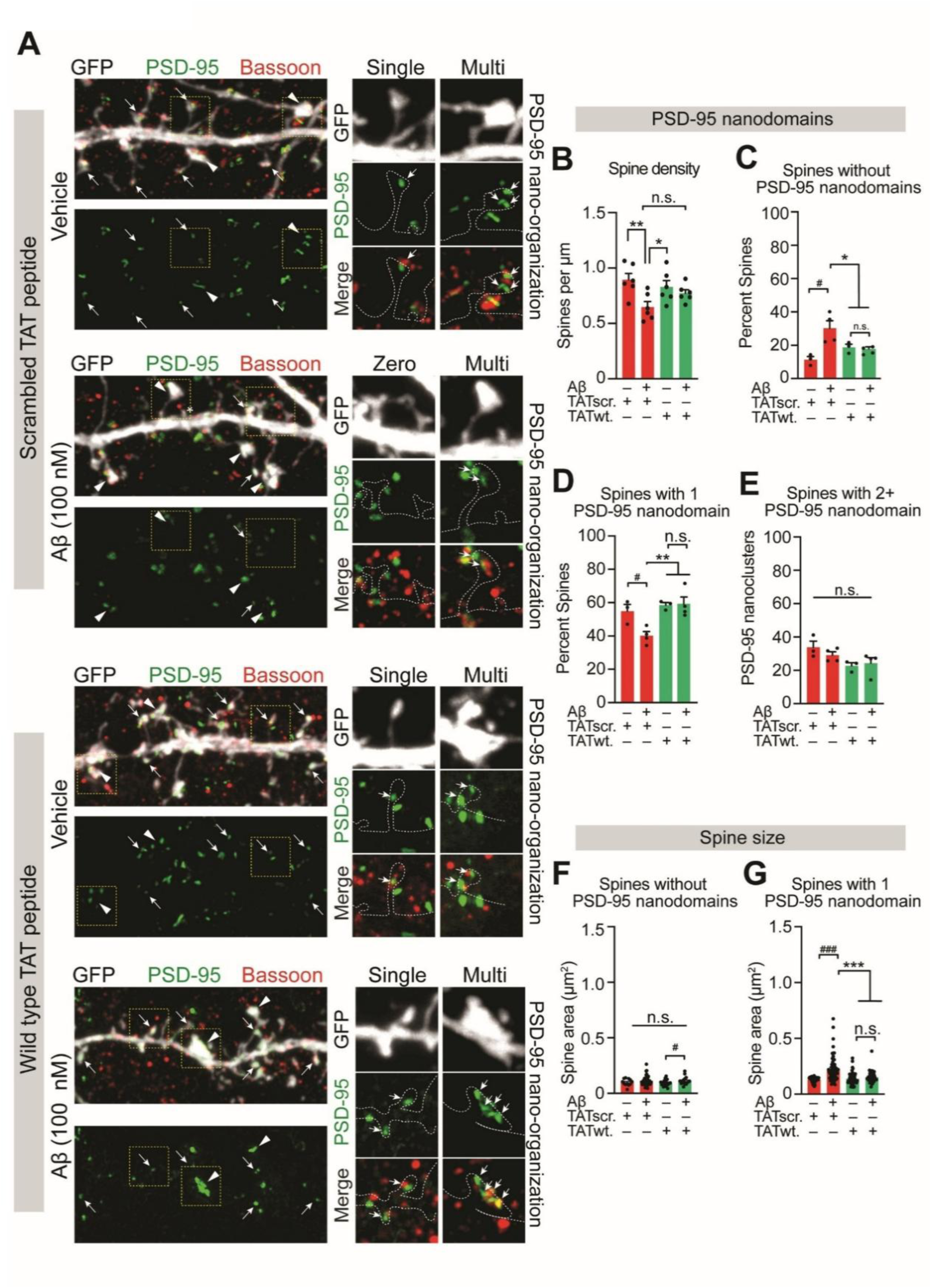
Displacement of β_2_AR from Ca_V_1.2 rescued Aβ-induced changes in spine nanoarchitecture. **A**: Representative tau-STED images of spines from DIV 21–25 cortical neurons transfected with GFP white) and labeled for PSD-95 (green) and Bassoon (red) after treatment with vehicle or 100 nM Aβ for 24 h with 10 µM of either control tat-Pep1923scr or active tat-Pep1923. Arrows indicate spines with one aligned PSD-95/Bassoon nanodomain and arrowheads spines with multiple nanodomains. Enlarged views of selected spines (yellow squares) with indicated PSD-95 nanodomain numbers are shown to the right. **B-E**: Overall spine density (**B**), and percentage of spines with 0, 1, or ≥2 PSD-95 nanodomains. Data points represent individual neurons (n 3/4 neurons for tat-Pep1923scr without/with Aβ and 3/4 neurons for tat-Pep1923 without/with Aβ). **F, G** Average size of spines with 0 or 1 PSD-95 nanodomain. Dots on graph represent individual spines (n for 0 nanodomains for tat-Pep1923scr: 16 without and 44 with Aβ; for tat-Pep1923: 21 without and 16 with Aβ; n for 1 nanodomain for tat-Pep1923scr: 52 without and 60 with Aβ; for tat-Pep1923: 61 without and 55 with Aβ). All data: mean ± SEM from at least 3 different neurons from 2 independent experiments (*p<0.05, **p<0.01, ***p<0.0001, ANOVA, Tukey’s post-hoc; ^#^p<0.05, ^###^p<0.0001, t-test).

**fig. S10.**
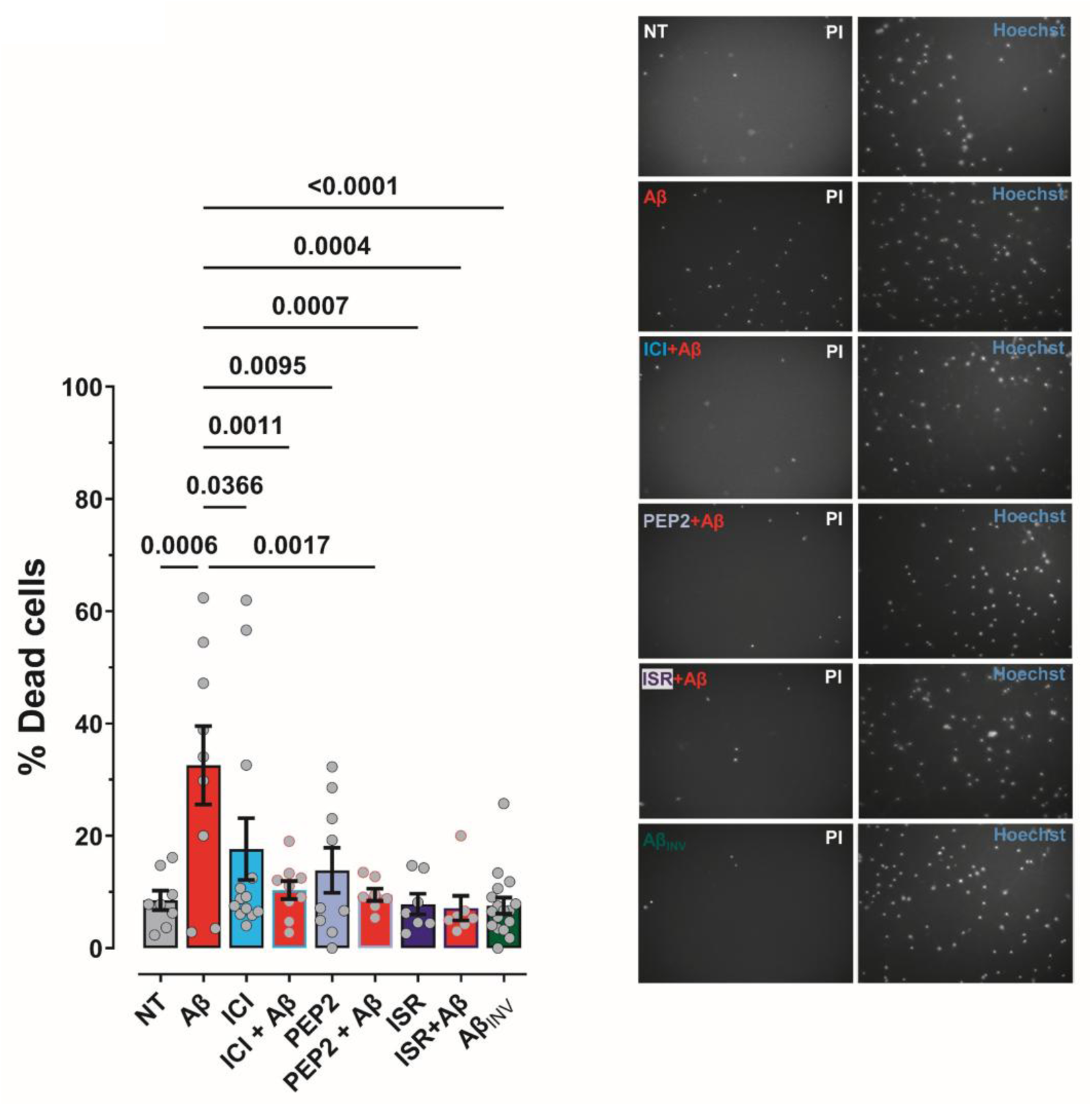
Blockade of β_2_AR signaling reduces Aβ-mediated neuronal death. HC at DIV 11-13 were treated with 1 μM Aβ (or Aβ_inv_) ± 10 μM isradipine (LTCC blocker) or 200 nM ICI 118551 (β_2_AR blocker) or 10 μM myrPep2 (myr-1923-1942 peptide) for 48 hours, followed by staining (10 min) with membrane impermeant propidium iodide (PI;10 μg/ml) to stain dead or dying neurons (with damaged plasma membrane) and membrane permeant Hoechst 33342 (10 μg/ml) to stain all neurons. Cell damage was reported as the percentage of PI positive cells over Hoechst-positive cells (n: 3-6 independent experiments; ** p<0.01, one-way ANOVA with Tukey’s post-hoc test; other than Aβ alone, no condition was significantly different from no treatment control).

